# Large-scale multivariate multi-ancestry Interaction analyses point towards different genetic mechanisms by population and exposure

**DOI:** 10.1101/562157

**Authors:** Vincent Laville, Timothy Majarian, Yun J Sung, Karen Schwander, Mary F Feitosa, Daniel Chasman, Amy R Bentley, Charles N Rotimi, L Adrienne Cupples, Paul S de Vries, Michael R Brown, Alanna C Morrison, Aldi T Kraja, Mike Province, C. Charles Gu, James Gauderman, DC Rao, Alisa Manning, Hugues Aschard, on behalf of the CHARGE Gene-Lifestyle Interactions Working Group

## Abstract

The *CHARGE Gene-Lifestyle Interactions Working Group* is a unique initiative formed to improve our understanding of the role and biological significance of gene-environment interactions in human traits and diseases. The consortium published several multi-ancestry genome-wide interaction studies (GWIS) involving up to 610,475 individuals for three lipids and four blood pressure traits while accounting for interaction effects with drinking and smoking exposures. Here we used GWIS summary statistics from these studies to decipher potential differences in genetic associations and GxE interactions across phenotype-exposure-population trios, and to derive new insights on the potential mechanistic underlying GxE through in-silico functional analyses. Our comparative analysis shows first that interaction effects likely contribute to the commonly reported ancestry-specific genetic effect in complex traits, and second, that some phenotype-exposures pairs are more likely to benefit from a greater detection power when accounting for interactions. It also highlighted a negligible correlation between main and interaction effects, providing material for future methodological development and biological discussions. We also estimated contributions to phenotypic variance, including in particular the genetic heritability conditional on the exposure, and heritability partitioned across a range of functional annotations and cell-types. In these analyses, we found multiple instances of heterogeneity of functional partitions between exposed and unexposed individuals, providing new evidence for likely exposure-specific genetic pathways. Finally, along this work we identified potential biases in methods used to jointly meta-analyses genetic and interaction effects. We performed a series of simulations to characterize these limitations and to provide the community with guideline for future GxE studies.

## Introduction

The precise role of gene-environment interactions (GxE) in complex human disease traits remains unclear. Although genome-wide GxE studies having been conducted for many phenotypes, the number of identified GxE is very small relative to the large number of genetic variants identified in traditional genome-wide association studies (GWAS). A number of issues related to the identification of GxE have been well described in the literature^1–3^, including in particular very low power^4^. As a result, the required sample size needed to detect GxE is substantially larger than for of marginal genetic effect. Moreover, few studies have explored potential differences in GxE across populations, or assessed the contribution of GxE to the variance of human phenotypes, or explored enrichment of GxE for specific functional mechanisms.

The Gene-Lifestyle Interactions Working Group^5^ (GLIWG) within the Cohorts for Heart and Aging Research in Genetic Epidemiology (CHARGE) is an international initiative that has the potential to address some of these challenges. It is a large-scale, multi-ancestry consortium that aims at systematically evaluating genome-wide gene-lifestyle interactions on cardiovascular disease related traits using genotypic data from up to 610,475 individuals. The consortium published a series of genome-wide single nucleotide polymorphism (SNP) by smoking and drinking interaction screenings focusing on four blood pressure phenotypes: diastolic blood pressure (DBP), systolic blood pressure (SBP), pulse pressure (PP), mean arterial pressure (MAP), and three lipid levels: triglycerides (TG), high-density lipoprotein cholesterol (HDL), and low-density lipoprotein cholesterol (LDL). For each pair of a phenotype and an exposure, a genome-wide interaction studies (GWIS) using the 1 degree of freedom (df) test for GxE interaction and the 2 df joint test of genetic and interaction effects^6^ has been conducted. The results from these analyses have been published in five papers: SNP-by-alcohol interaction^7^ and SNP-by-smoking interaction^8,9^ on blood pressure, and SNP-by-alcohol interaction^10^ and SNP-by-smoking interaction on lipids^11^.

Here we first synthesize the GWIS results for all phenotype-exposure combinations. We highlight the importance of our large-scale initiative, providing evidence that interacting variants might differ by genetic ancestry, and show that accounting for GxE can help discovering new loci, especially for certain phenotype-exposure pairs. We then performed a series of analyses comparing interaction effects against both genetic main effects estimated in our studies and marginal effects from previous GWAS. Contrary to a commonly assumed hypothesis^12^, we found only negligible correlation between the interaction and marginal effect, highlighting additional challenges for future GxE interactions studies. Estimated variance explained by main and interaction effect for the outcomes under study also showed that in general, interactions explain a very small amount of phenotypic variance on top of the marginal genetic effect for these traits. However, these limitations were balanced by heritability analyses. Partitioning the genetic variance in exposed and unexposed individuals separately, using both functional and cell type annotations, we found differential enrichment between the two groups in multiple instances. This suggests GxE might still play an important role in these phenotypes, with some exposures potentially triggering new molecular mechanisms or reducing the contribution of pathways involved in unexposed individuals.

## Material and Methods

### Phenotypes and exposures

We considered four blood pressure phenotypes (DBP, SBP, PP, MAP), and three lipids levels (TG, HDL, LDL). DBP and SBP were derived as the average over multiple measurements performed at resting or sitting positions. PP and MAP were derived as the difference between SBP and DBP, and the sum of two-thirds of DBP and one-third of SBP, respectively. In all cohorts, HDL and TG were directly assayed, while LDL was either directly assayed or estimated using the Friedewald equation^13^: LDL = TC − HDL − (TG/5). Both HDL and TG were natural log transformed, while LDL was not transformed. Additional details of the phenotype transformation have been published here^5^.

Two binary smoking exposures, *current smoking* and *ever smoking*, were considered and measured similarly across all smoking GWIS. The *current smoking* variable was coded as 1 if the subject smoked regularly in past year and as 0 otherwise. *Ever smoking* status was coded as 1 if the subject smoked at least 100 cigarettes during his/her lifetime and 0 otherwise. For alcohol consumption, two binary variables were considered, referred further as *current drinking* and *drinking habit*. For both blood pressure and lipid traits, the former exposure was defined similarly for all studies, corresponding to any recurrent drinking behavior. The *drinking habits* exposure was defined differently across publications. For lipids phenotypes, the variable was coded as 1 for the subset of current drinkers having at least two drinks per week and 0 to for everyone else (i.e. the no drinkers and those drinking less than two drinks per week)^10^. The blood pressure GWIS used instead a “low *versus* heavy drinking”, where the variable was coded 1 for individuals having at least 8 glasses per week, and 0 for individuals with less than 8 glasses per week, while all non-drinkers were removed^7^.

Generally, the use of categories for the exposures was necessary for harmonizing data from the large number of studies, especially for alcohol consumption. Additional details on the assessment of the exposure and phenotypes are provided in the corresponding publications.

### Data pre-processing

All studies conducted a two-stage approach. In stage 1 (referred as *Discovery*), a standard GWIS was performed using up to 18 million genetic variants. In stage 2 (referred as *Replication*), only a subset of variants with a *p*-value below a certain threshold (P<10^−6^ or P<10^−5^) at stage 1 were further considered. More details can be found in the corresponding publications^7,8,10,11^. For each outcome-exposure, we had access to complete meta-analysis summary statistics of both the discovery and the replication stages for populations of four different ancestries (European, African, Asian and Hispanic) after quality control filtering. To ensure a fair comparison, we re-processed all results using the same pipeline. In the discovery stage, we excluded SNPs with a MAF below 1% and with significant (*P* < 10^−6^) heterogeneous effects across individual cohorts. SNPs present in only one ancestry were excluded from trans-ancestries analyses. Trans-ancestry summary statistics in the replication stage were filtered similarly to the discovery stage. Finally, we computed meta-analyses results for the combined analyses (discovery stage + replication stage) in each individual ancestry and trans-ancestry. For each ancestry and each phenotype-exposure combination, only SNPs included in both stages were retained in the final combined dataset. All meta-analyses were computed using the METAL software^14^.

### Identification of independent signals and loci

We report genome-wide significant variants in the combined meta-analyses (*P* < 5×10^−8^) for each outcome-exposure and in each ancestry. Independent signals were defined using the clumping framework from the PLINK software^15^, using a linkage disequilibrium (LD) threshold of 0.2 and a maximum physical distance from the lead SNP (i.e. the most associated variant) of ± 500 kb. The LD was derived using 1000 Genomes Project^16^ individuals as a reference panel while accounting for ancestry. We used the EUR, AFR, combined EAS-SAS and AMR samples as proxies for the individuals from European, African, Asian and Hispanic ancestries, respectively. For the trans-ancestry analyses, we built our reference panel by merging all these populations. In some tables and figures, we also grouped independent association signals into loci by clustering SNPs located less than 500 kb upstream or downstream from the lead SNP. Note that when deriving shared associated loci across studies (e.g. across ancestries), we merged loci that overlap, resulting in total counts sometimes slightly lower than the expected total count.

### Interaction effect conditional on marginal effect

We assessed potential enrichment for interactions effects for SNPs displaying marginal genetic association. To ensure independence between our interaction effect GWIS and the marginal GWAS, we used summary statistics from previous studies on blood pressure traits^17–19^ and lipid traits^20–23^. Here, we considered only individuals of European ancestry, in order to maximize the sample size while limiting potential issues due to genetic heterogeneity, where the top variants might differ across populations. Moreover, to avoid enrichment driven by a single locus, we performed a clumping of the GWAS of marginal genetic effect with PLINK^24^, so that all candidate SNPs considered are independent from each other. We first derived the proportion of interaction effect nominally significant at type I error rate (alpha) threshold of 0.05 among successive bins of SNPs selected based on their marginal association. For the last bin, including only SNPs previously identified at genome-wide significance level, we also performed three complementary approaches to test jointly interaction effects^4^ at those variants: an omnibus test, an unweighted genetic risk score (uGRS) test, and a weighted genetic risk score (wGRS) (see **Supplementary Note**).

### Variance explained and heritability

We first estimated the fraction of phenotypic variance explained by top SNPs, decomposed into main effects, interaction effects and those effects jointly using the R package *VarExp*^25^ (see **Supplementary Note**). The analysis was conducted for each ancestry and each phenotype-exposure combination separately, using only genome-wide significant SNPs in the combined meta-analyses for either the 2df or the 1df test for the given trio (exposure-phenotype-ancestry). For simplicity, we clustered SNPs into loci of 1Mb (500kb from the top SNP upstream and downstream) and computed the variance explained using only top SNPs (with the lowest p-value) for all loci. Also, because of potentially biased estimations of the interaction effect sizes using the 2df framework but not for the main genetic effect size, we used the genetic main effect size estimates from the joint framework and the interaction effect sizes computed using the standard 1df meta-analyses for the interaction test.

For each project, we also aimed at assessing potential differences in heritability across exposure-specific strata. Again, to avoid genetic heterogeneity issues, we focused on European ancestry samples only. We computed the genetic heritability in the whole sample and in exposure-specific strata (i.e. in unexposed and exposed individuals separately) for each trait and exposure combination using the *LDscore* approach^26^. We used the pre-computed *LDscores* relative to European ancestry samples provided with software. When unavailable from the original studies, we derived the summary statistics of the genetic marginal effect in the whole sample and in unexposed and exposed individuals from the interaction model using a tool we recently developed^27^.

### Stratified heritability

For each exposure stratum, genetic heritability was further partitioned by both cell type-specific and general annotations^28^. As for the overall heritability, these analyses focused on European ancestry individuals only. We used two distinct sets of annotations: *baseline* and *GenoSkyline+*. The *baseline* annotations encompass 53 tissue-agnostic, general functional annotations. *GenoSkyline+* is a recently proposed annotation set integrating a rich collection of epigenomic data from the Roadmap Epigenomics Project^29^. Additional details on the annotations are provided in the **Supplementary Note**. Enrichment and annotation-specific heritability were compared across exposure strata for each trait. When assessing the significance of the enrichment, we used a Bonferroni corrected significance threshold of P < 0.000277. We further quantified enrichment for tissue-specific heritability following Finucane et al.^30^, where results from cell-specific annotations based on gene expression data were gathered into tissue-specific classes. Except when specified otherwise, enrichment analyses compared median enrichment between exposure strata, avoiding comparison of significance which would be biased by differences in sample size.

### Simulation study

We compared the performances of meta-analysis strategies for estimating and testing the main and interaction effects across multiple cohorts. The first strategy (1df framework) uses the effect estimate and standard error of the parameter of interest (either the main or interaction term) from each individual cohort and then performs a standard 1 degree of freedom inverse-variance weighted meta-analysis. The second strategy (2 df framework) performs first a meta-analysis of both parameters jointly, using not only single cohort effect estimates and standard errors, but also their covariance^14,31^. It then uses the effect estimates from the previous step to perform a standard *Wald* test of each parameter separately.

We conducted several simulations, all including two cohorts for clarity. The first focused on understanding differences we observed for the interaction effect between the two frameworks on real data. We generated 20,000 genotypes per cohort, a binary exposure and a phenotype as a linear combination of a main genetic effect, an interaction effect, or both. These simulations aimed at assessing the impact of heterogeneity across cohorts, when varying the MAF of the SNPs, the proportion of exposed individuals in the two cohorts, and the effect sizes of the main genetic effect and of the interaction effect. For each replicate and each scenario, we perform a linear regression in each cohort and applied the two aforementioned frameworks to test for the interaction between the SNP and the exposure. The second simulation broadened the scope of the assessment, and compared estimated coefficient and corresponding chi-squared test for both the interaction and main genetic effect terms for the two frameworks. Here, we simulated a series of 1,000 replicates using either a binary outcome or a continuous outcome and a single SNP, while varying all parameters (distribution of the exposure, MAF, presence and size of the genetic and interaction effects, sample size of each cohort) at random and independently between the two cohorts. Finally, the last simulation uses a similar framework but focuses on comparing the power and robustness of the 2df joint test of main and interaction effect, as compared to marginal genetic effect model.

## Results

### Overview

We focused on three lipid and four blood pressure phenotypes, each examining GxE interaction with two smoking and two alcohol exposures, for a total of 28 GWIS (**Table 1**). All outcome-exposure pairs considered were analyzed using a two-stage approach involving up to 610,475 individuals. In stage 1, genome-wide interaction analysis was performed in up to 29 cohorts with a total of up to 149,684 individuals from multiple ancestries: European-Ancestry (EA), African-Ancestry (AA), Asian-Ancestry (ASA), and Hispanic-Ancestry (HA). In stage 2, involving up to an additional 71 studies with 460,791 individuals, also from multiple ancestries, studies focused on the replication of a subset of variants from stage 1 with a *p*-value threshold of 1.0 × 10^−6^ achieved by either the 1df or the 2 df test. Note that the total sample size (discovery + replication) varied substantially across the trait analyzed, with an average of 311K for lipids and 457K for blood pressure traits. To ensure a fair comparison across all analyses, we re-processed all GWIS summary results using the same pipeline. Stage 1 quantile-quantile (QQ) plots for both the 1df and the 2df test are presented in **Figure S1**, and frequency of the exposure are presented in **Figure S2** and **Table S1**. Finally, note that the primary association results from the original studies and our analyses are highly concordant, but minor differences might exist because of slight differences in the analysis pipeline.

**Table 1.**
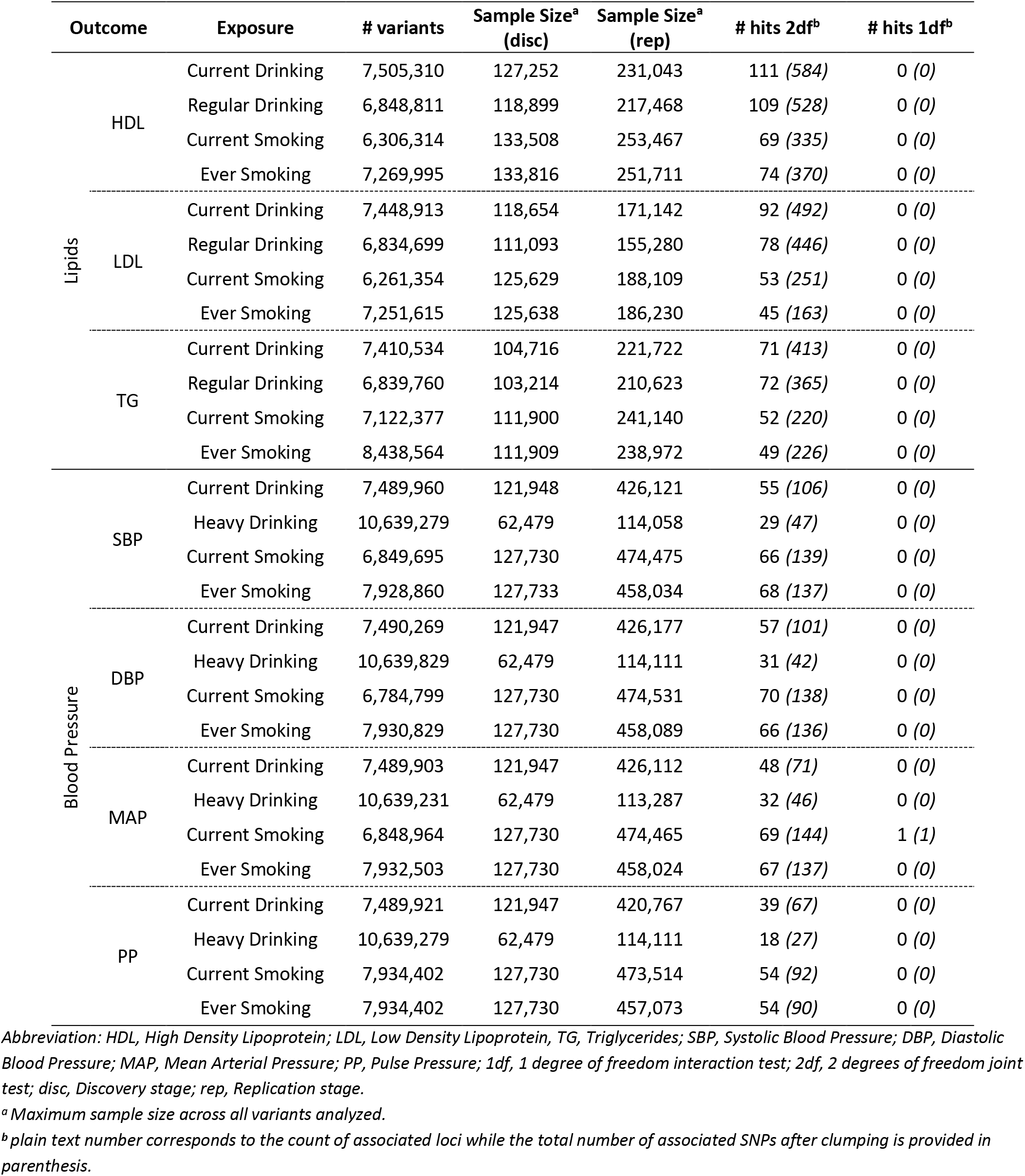
Summary of trans-ancestry GWIS results for 2df joint and 1df interaction tests.

### Summary of 2df results

The 2df test identified a large number of variants in both the trans-ancestry (**Table 1**) and ancestry-specific (**Table S2**) meta-analysis. After clumping SNPs based on their pairwise linkage disequilibrium, the 2df trans-ancestry analyses identified a total of 5,913 association signals, reduced to 1,698 associations when aggregating neighboring SNPs into loci (see ****Material and Methods****). A total of 54% of loci (*N*=926) harbored a single independent association signal (**Figure S3**). For the other loci, the number of potentially additional signals equaled 3 on average with a maximum of 71. Importantly, many loci overlapped across the exposures tested. For example, there were 108 and 103 loci identified for HDL when including interaction between current drinking and drinking habits, respectively. However, 92 of those loci overlap between the two analyses. Further merging all overlapping loci identified by different exposure scans, our studies found a total of 112, 98, 77 loci for HDL, LDL, TG, and 74, 75, 75 and 59 loci for SBP, DBP, MAP and PP. On average, 13% of the loci were identified by a single exposure scan, while 41% were identified by all four exposure association studies for each phenotype (**Figure 1a**).

**Figure 1.**
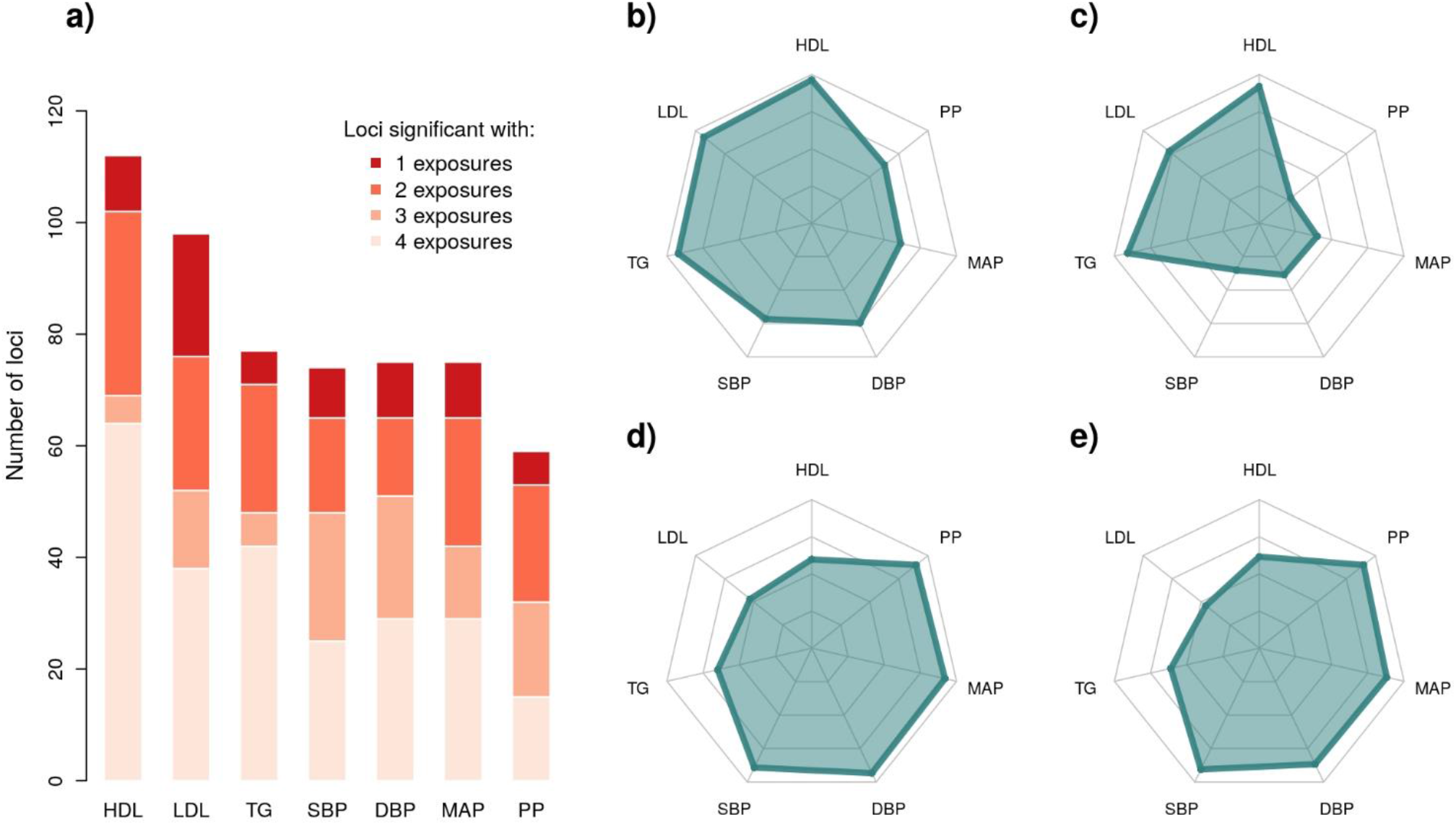
Loci identified by the trans-ancestry 2df joint test across the four exposures. We assessed the relative performance of the trans-ancestry joint 2df test across the four exposures. Panel **a)** shows overlapping loci for the 2df test across the four exposures. We further decomposed these results by exposure, for current drinking (**b**), drinking habit (**c**), current smoking (**d**), ever smoking (**e**). The corresponding radar plots show the proportion (from 0% to 100%) of the total number of loci identified for that phenotype.

When stratifying results by exposure, accounting for interaction with drinking tended to identify more lipids associations, while accounting for interaction with smoking identified more associations for blood pressure phenotypes (**Figure 1b-e**). Looking at cross-phenotypes results, GWIS accounting for current drinking and drinking habits captured 81% and 61% of all loci, respectively, and current smoking and ever smoking scans identified 75% and 72% of all loci respectively. Note that the lower number of signals for drinking habits is likely partly explained by the smaller sample size used for that exposure (307K on average versus 440K for the other exposures), especially for the BP GWIS that used a different definition for drinking habits (see ***Material and Methods***). Nevertheless, to understand the differences observed across other exposures, we used the HDL results as a case study. First, we noticed that the 2df chi-squared test from overlapping loci across the four exposure scans were highly correlated (**Figure S4a**). This is expected, as most studies have approximately the same sample size at discovery and replication stages, and assuming the contribution of the interaction effect is limited. Conversely, we noticed a larger mean interaction effect chi-square at those same loci for the drinking exposures as compared to the smoking exposures, whatever the framework used to derive the interaction chi-squared test (see the last result sections), suggesting the higher detection rate is at least partly explained by a contribution of the interaction effect (**Figure S4b**).

An important novelty of the Gene-Lifestyle Interactions Working Group is the inclusion of a large proportion of non-European ancestry individuals. More precisely, over the two stages, 63% (*N*=380,612) were of European, 27% (*N*=162,370) of Asian, 6% (*N*=34,901) of African and 4% (*N*=22,334) of Hispanic ancestries. For the 2df test, the total number of significant associations per ancestry was globally proportional to the available sample size (**Table S1**). Merging results from all phenotype-exposure pairs, there was 1,285, 383, 135, and 148 phenotype-variants associations identified after clumping by this approach in EA, ASA, AA, and HA, respectively. Deriving the overlap across ancestries for those associations, we found that the vast majority of ASA and HA associations were also identified by the larger EA studies (**Figure 2a**). Conversely, 32% (43 out of 135) of the associations identified in AA were exclusively identified in this population. These association mostly implied variants monomorphic in all ancestries except the African ancestry. The trans-ancestry analysis identified 1276 (94%) of all ancestry-specific associations, while uncovering an additional 148 associations. All associations missed in the trans-ancestry analyses were found in a single ancestry from ASA (N=6), AA (N=36), EA (N=41), and HA (N=1). To account for sample size differences and assess whether top variants were consistent across populations, we also extracted for each ancestry-specific signal the *p*-value at the same top SNP for the other ancestries from stage 1, and assumed replication if that *p*-value was smaller than 0.05. **Figure 2b** shows the overlap over all phenotypes and per phenotype is modest, which suggests enrichment for ancestry-specific variants in most populations and in African ancestry in particular.

**Figure 2.**
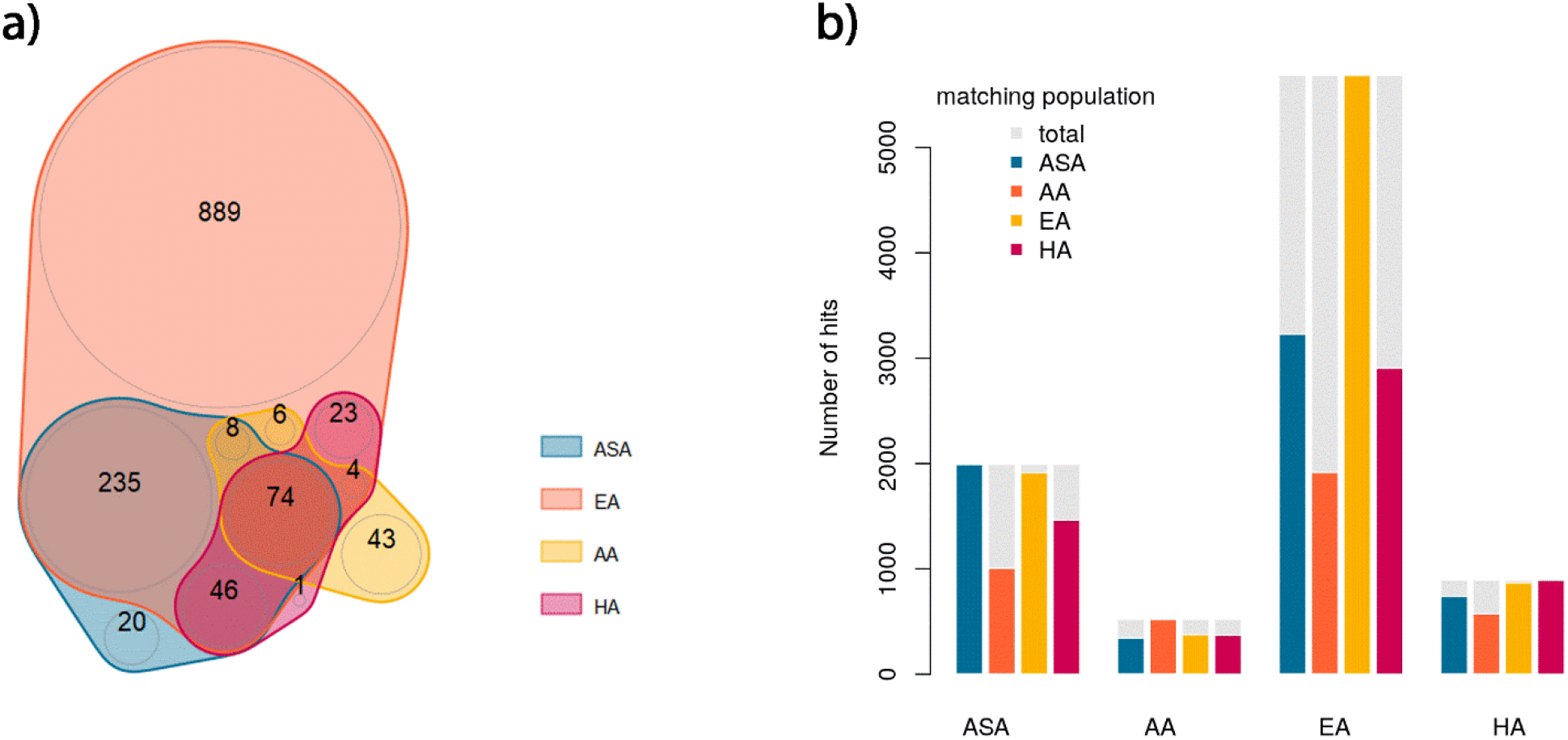
Overlapping associations for the 2df test across ancestries. We derived the overlap in association signal for the joint 2df test of main and interaction effects across the four ancestries: Asian (ASA), African American (AA), European (EA), and Hispanic (HA). Panel **a)** shows a Venn diagram focusing only on loci found at genome-wide significance level after the meta-analysis of stage 1 and 2. In panel **b)** we extracted genome-wide significant and independent SNPs per ancestry (*i.e.* reference population) after the meta-analysis of stage 1 and 2, and extracted the *p*-value for those SNPs in other population (*i.e.* the matching population) from stage 1. The barplot shows for each reference population, the proportion of SNPs in the matching population that achieve a *p*-value below 0.05.

### Summary of 1df results

Despite the reasonably large sample size available in our studies, we found only one significant interaction across the 28 trans-ancestry GWIS when combining discovery and replication. Nevertheless, we attempted to assess a potential GxE contribution by decomposing the signal from the significant 2df results. For each phenotype-exposure-ancestry trio we derived the number of SNPs inducing an enhanced genetic effect in exposed individuals (when main and interaction have the same direction) and those inducing a reduced genetic effect (when main and interaction have opposite signs). As showed in **Figure S5**, main and interaction tended to be distributed at random among those SNPs, although we did observe a slight enrichment for significant differences, with 14 out of 91 trios showing nominally significant (*P*<0.05) disequilibrium in concordant versus discordant effects. Four of them, all in trans-ancestry analyses, remain significant after correction for multiple testing: LDL showed larger genetic effect in both current and ever smokers, DBP showed larger genetic effect in current drinkers, and SBP showed smaller genetic effect among ever smokers. Among sets of variants displaying interaction effects discordant with main genetic effect, we also searched for those inducing an opposite effect between exposed and unexposed individuals. Although it is expected that the 2df test is supposed to overperform the marginal test, there was only 41 such associations (0.4% of all associations).

Finally, in ancestry-specific meta-analysis, the 1df interaction test identified 8 loci reaching genome-wide significance, all observed in the African ancestry population (**Table S3**). They involved smoking exposure only and are associated with both lipids and blood pressure traits. Four were also detected at genome-wide significance with the 2df test in the African ancestry population, while the remaining four were relatively close to genome-wide significance with that test (*P* between 1.9×10^−6^ and 6.3×10^−8^). In line with the results from the 2df test (**Figure 2**), African ancestry appears to be enriched for ancestry-specific associations.

### Comparison against marginal effect screening

We retrieved from the literature loci exhibiting significant marginal genetic effect on blood pressure traits^17–19^ and lipid traits^22,32^, and compared those associations against both 1df and 2df tests from our stage 1 analysis (as some SNPs were not available at stage 2). Description of these references are provided in **Table S4**, and the list of SNPs used in **Table S5**. Overall, the 2df screenings identified 167 novel loci-outcome associations, where loci are defined as the genetic region ±500kb around the top associated variant. Among the 647 associations retrieved from the literature, 302 were also found in our studies, while 345 associations were not replicated at genome-wide significance level (note that this is a stringent comparison as some of them might be captured at stage 2). Most of the new association results for lipids were identified when accounting for interaction with drinking exposures, while the majority of new blood pressure associations were identified when accounting for interaction with smoking exposures (**Table 2**). For example, 86% (N=18) of the 21 new associations with HDL were found in the gene-by-current drinking GWIS, when only 52% (N=511) were identified in the gene-by-current smoking GWIS. Conversely 65% (N=17) of the 26 new associations with DBP were found in the gene-by-ever smoking GWIS, while 15% (N=4) were identified in the gene-by-drinking habits GWIS.

**Table 2.** Association signal overlap between the 2df test (accounting for interactions) and previous GWAS of marginal genetic effect.

| Phenotype | Overall |  |  | CHARGE only, per exposure |  |  |  |
| --- | --- | --- | --- | --- | --- | --- | --- |
|  | *External GWAS only* | *both* | *CHARGE only* | *Current Drinking* | *Drinking Habits* | *Current Smoking* | *Ever Smoking* |
| **HDL** | 70 | 83 | 21 | 18 | 17 | 11 | 11 |
| **LDL** | 60 | 64 | 29 | 24 | 16 | 5 | 6 |
| **TG** | 91 | 58 | 14 | 9 | 11 | 5 | 4 |
| **SBP** | 37 | 43 | 29 | 16 | 2 | 21 | 22 |
| **DBP** | 47 | 46 | 26 | 16 | 4 | 22 | 17 |
| **MAP** | - | - | - | - | - | - | - |
| **PP** | 40 | 8 | 48 | 33 | 15 | 45 | 44 |
| **All** | 345 | 302 | 167 | 116 | 65 | 109 | 104 |

Abbreviation: HDL, High Density Lipoprotein; LDL, Low Density Lipoprotein, TG, Triglycerides; SBP, Systolic Blood Pressure; DBP, Diastolic Blood Pressure; MAP, Mean Arterial Pressure; PP, Pulse Pressure; 1df, 1 degree of freedom interaction test; 2df, 2 degrees of freedom joint test; disc, Discovery stage; rep, Replication stage.

We next assessed potential enrichment for interaction effects across variants previously identified in these marginal effect GWAS. The distribution of interaction effects at those variants did not indicate any clear trend (**Figure S6**) and the joint test of all single SNP^33^ did not find any enrichment for interaction effect among these variants (**Table S6**). The smallest single SNP *p*-value was observed for rs1260326 (*P* = 3.3e-6), a missense variant in *GCKR*. Interestingly, *GCKR* has been previously found associated with alcohol consumption^34,35^, and interaction between variants in *GCKR* and alcohol consumption have been reported for gout disease^36^. Besides assessing interaction effect at genome-wide significant variants, we also explored potential enrichment for interaction at non-significant SNPs. Such enrichment would be of particular interest to increase power of GxE test through 2-step approaches^12,37,38^ (see for example **Figure S7**). The most common 2-step approach consists of filtering out SNPs displaying a marginal genetic *p*-value larger than a given α1 significance threshold. To assess for the potential of this strategy in our data, we quantified the enrichment of nominally significant variants (*i.e. P* < 0.05) for GxE interaction effect while varying α1 between 0.1 and 10^−6^ applied to the aformentioned external marginal GWAS summary statistics. Overall, there was no clear enrichment in our data (**Figure 3**), although some phenotype-exposure pairs show a slight increase in the proportion of significant GxE interaction, including in particular TG and drinking habits (11% of the SNPs against the 5% expected for α_1_=10^−5^). Note that the absence of enrichment for other phenotype-exposure pairs does not rule out the relevance of this strategy, but suggests that alternative metrics might be used to select candidates.

**Figure 3.**
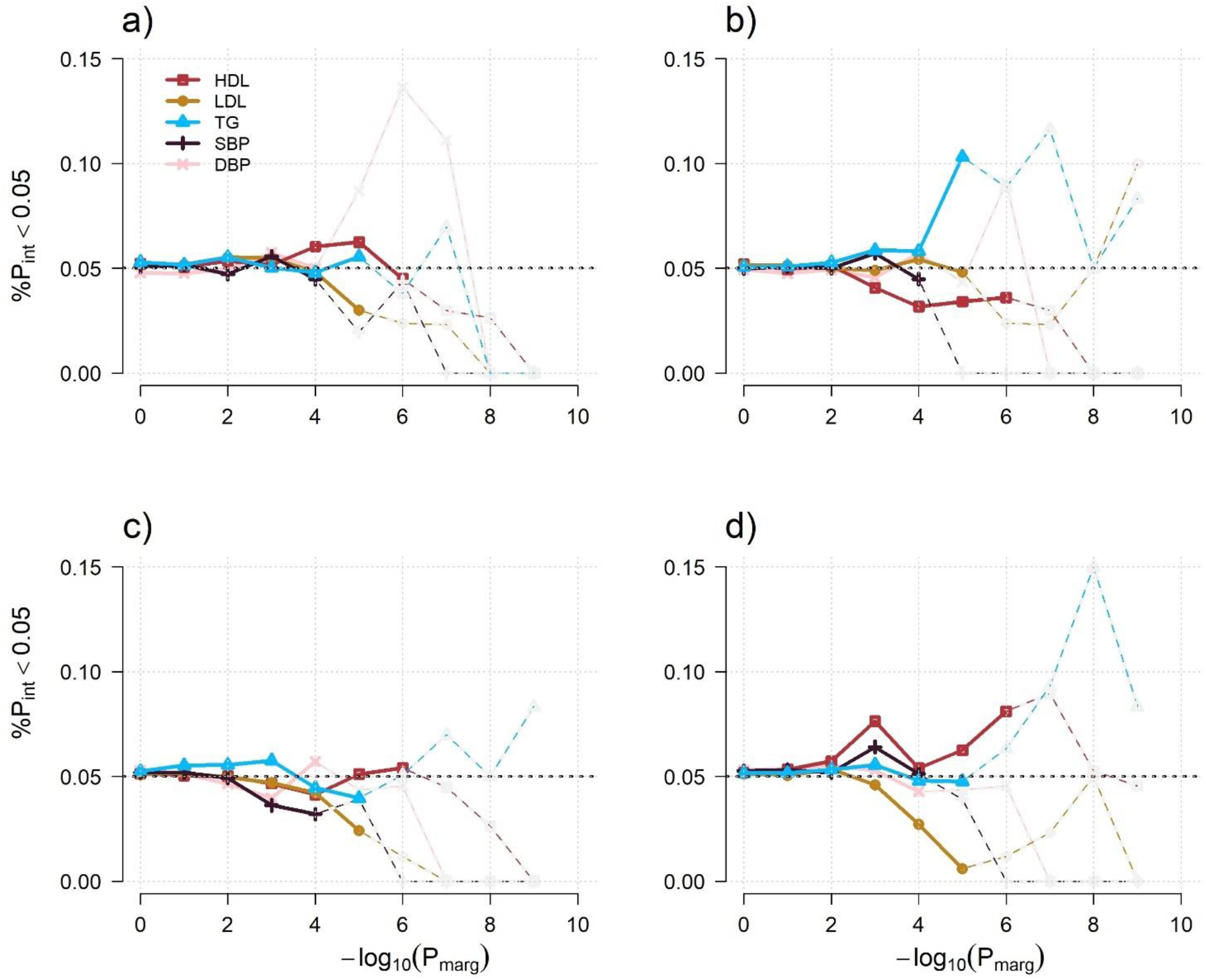
Potential power for 2-step approach. We plotted for each environmental exposure, current drinking (a), drinking habits (b), current smoking (c) and ever smoking (d), the proportion of independent SNPs displaying an interaction *p*-value (P_int_) below 0.05 in CHARGE across bins of variants selected from an independent marginal effect GWAS. Those bins were defined as sets of independent variants with *p*-value for marginal genetic effect (P_marg_) lower than a given threshold. All analyses used only GWIS results from European ancestry individuals. Under the null hypothesis of no correlation, we expect the proportion to be close to 0.05 (the black dashed line), independent of the threshold for P_marg_. Each of the five phenotypes are represented by a plain color line. When bins of SNPs harbor less than 100 variants (for low p-value threshold), proportion of P_int_ < 0.05 are indicated by dashed lines.

### Variance explained

We first used VarExp, a tool we recently developed^25^, to estimate the variance explained by marginal genetic effects, the joint genetic and GxE interaction effects, and the interaction effect only at the top genome-wide significant variants in each locus for each phenotype-exposure-ancestry analysis (**Table S7**). Overall, marginal genetic effects explained between 0.09% and 8.72% of the total phenotypic variance with an average of 3.59%. On the other hand, the interaction terms explained between 0% and 0.41% of the phenotypic variance. The largest amount of variance explained was observed for lipids traits, (e.g. average of 4.47% for the 2df, as compared to 0.81% for blood pressure phenotypes). Overall, we did not observe any major difference in the fraction of variance explained by the main genetic effect or jointly by the main genetic effect and the interaction effect across populations. However, we note a larger fraction of variance explained in the European ancestry samples than in other populations, with greater differences observed in lipids phenotypes and drinking exposures (7.11% of explained variance in individuals from European ancestry versus 5.11% in other populations on average). As expected, the fraction of variance explained by the interaction effects only were relatively small for all phenotype-exposure-ancestry trios. However, this fraction was slightly higher in the African ancestry population (0.15%) than in other ancestries (around 0.04%), in agreement with the higher number of significant interactions identified in the African samples.

Second, we estimated potential changes in the heritability of the three lipids and two blood pressures (DBP and SBP) traits across all individuals and in strata defined by exposure, using the *LDscore* approach^26^ applied to summary statistics from the analyses performed in the European ancestry population (**Figure 4**). We first found multiple phenotype-exposure pairs where heritability was significantly different between the exposed and unexposed groups. However, in most of the latter cases we noticed unexpectedly large values for the ratio measuring the proportion of the inflation in the mean chi-squared statistic that the LD Score regression ascribes to causes other than polygenic heritability. The maximum of this ratio equals 0.91 for SBP and drinking habits when the guideline suggests it should be close to zero. Such large ratio may indicate a partial mismatch between sample and reference for the LD Score or a potential model misspecification (e.g. when low LD variants have slightly higher heritability per SNP). We therefore performed a sensitivity analysis, re-deriving the heritability after filtering out SNPs based on their *p*-value for heterogeneity in the meta-analysis and selected the most reliable estimate (see **Figure S8** and **Supplementary Notes**). While some of the initially observed differences disappeared, most of the trends remained. In particular, heritability among exposed was on average smaller than among non-exposed for current smoking (h^2^=0.06 and h^2^=0.11, respectively) and for drinking habits (h^2^=0.15 and h^2^=0.12, respectively). Conversely, heritability was on average larger for current drinkers than non-current drinkers (h^2^=0.12 and h^2^=0.15, respectively).

**Figure 4.**
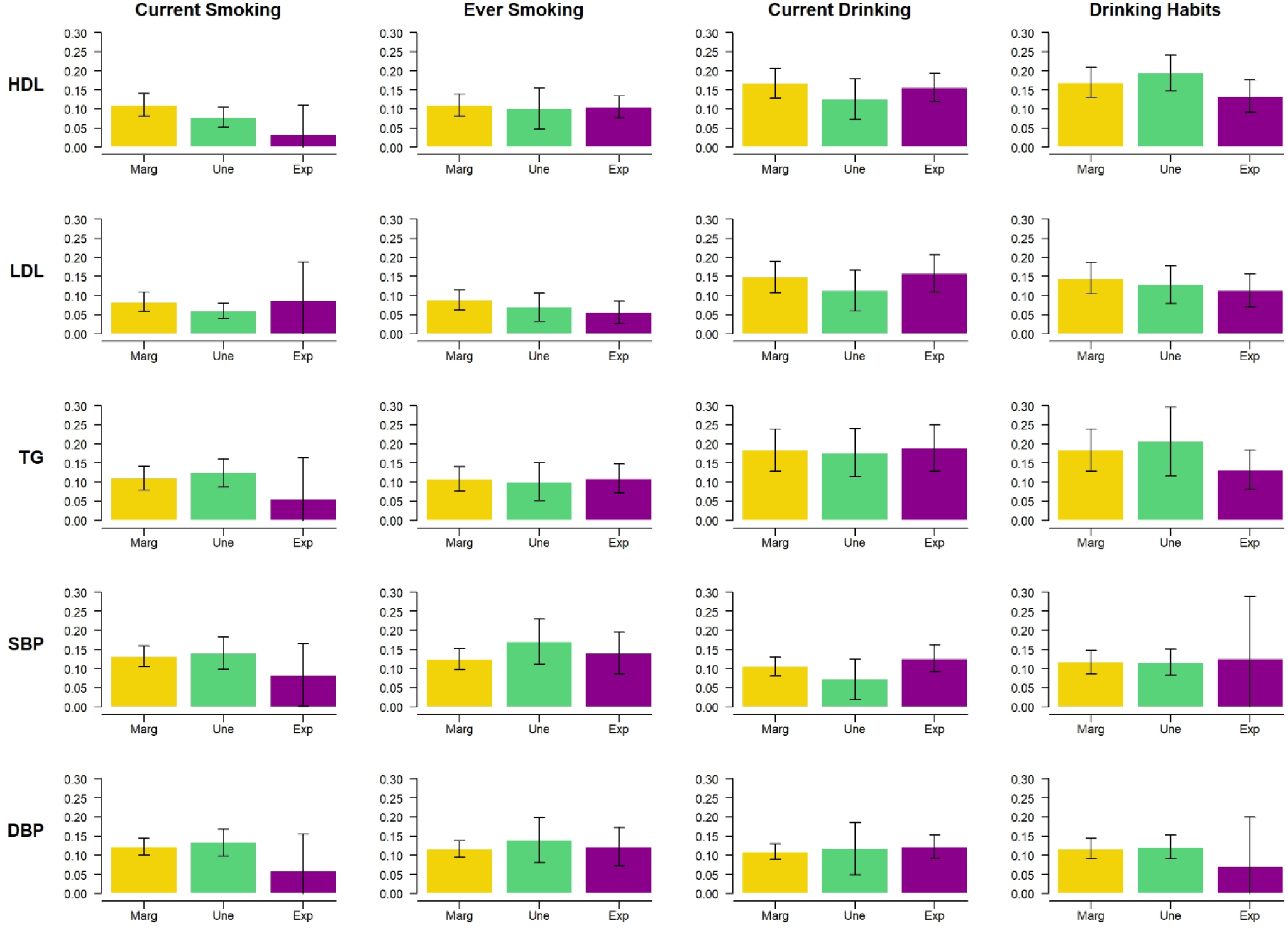
Heritability by exposure group. Heritability of the three lipids and two blood pressure phenotypes (DBP and SBP) derived using the *LDscore* applied to summary statistics from the European ancestry samples meta-analysis. Heritability was derived for all individuals (*Marg*, yellow bar) and for subset of unexposed (*Une*, teal bar) and exposed (*s*, purple bar) individuals. Vertical dark lines represent the 95% confidence intervals.

### Differential pathways across exposures

To explore further differences in genetic effect across exposure strata, we performed a second heritability analysis, partitioning genetic contribution by functional annotations^28,30^. We first considered baseline annotations provided with the *LDscore* package and the *GenoSkyline*^39^ annotation set, a cell-type specific annotation database derived mainly from the Roadmap Epigenomics^29^ (**Figures S9-13**). Because of the relatively modest sample size in some strata for such analyses (N equals 12,578 in the smallest strata, see **Table S1**), we focused on the distribution of estimated enrichment coefficient between exposed and unexposed. The majority of phenotype-exposure pairs show a similar enrichment pattern (**Figure 5**). For example, the enrichment estimates were highly correlated for drinking habits exposure and lipids (0.75, 0.59 and 0.39 for HDL, LDL and TG, respectively), suggesting that potential GxE interactions for those phenotypes do not involve new pathways. Conversely, LDL show substantial variability in enrichment for the three other exposures (correlation equals 0.10, 0.22, and 0.17, for current drinking, current smoking and ever smoking), suggesting those exposures might activate new genetic pathways while reducing the effect of genetic variants involved in unexposed populations. We also noted substantial variability for the phenotypes-exposure pairs showing the largest differences in heritability (lipids and current smoking, and BP and drinking habits, see **Figure 4**). However, part of that variability might be due to the reduced sample size in one of the two strata, thus making interpretation challenging.

**Figure 5.**
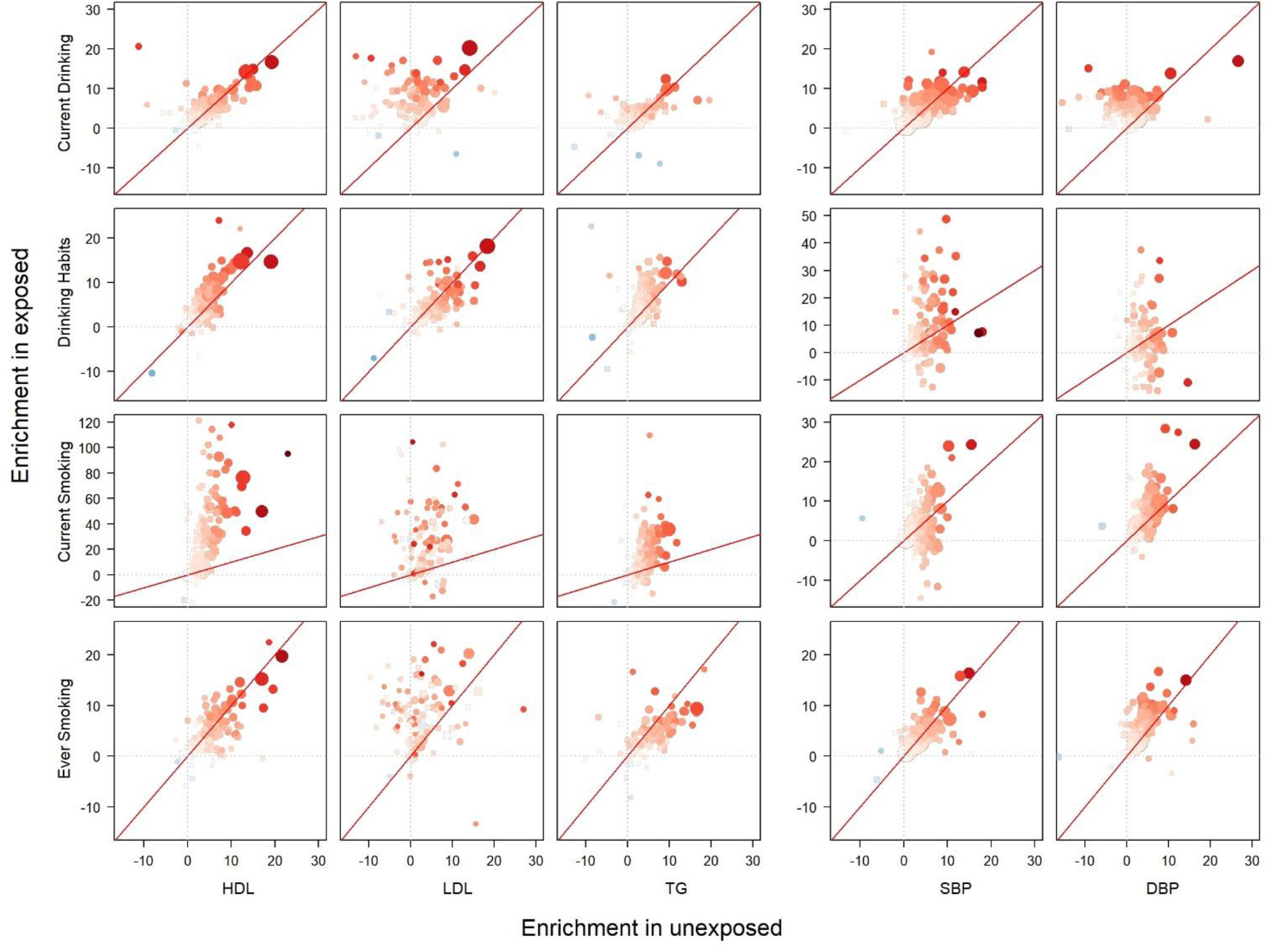
Functional enrichment across baseline and GenoSkyline+ annotations. Partitioned heritability for each lipid-exposure in European ancestry individuals across 180 functional annotations. Enrichment among unexposed individuals (X axis) is plotted against enrichment among exposed individuals (Y axis). Color and size of each data point indicates enrichment and significance of the enrichment in the total sample including exposed and unexposed individuals.

We next investigated whether exposures tend to display systematic enrichment in specific tissues^30^. For each phenotype, heritability was stratified based on annotation from 205 cell-types linked to 9 tissues (adipose, blood/immune, cardiovascular, central nervous system, digestive, endocrine, liver, musculoskeletal/connective, and other), in unexposed and exposed individuals separately. Again, because of unbalanced sample size between strata, we focused on the relative differences in median enrichment between exposed and unexposed by tissue, while significance was accounted for after merging strata. Detailed results per phenotypes are presented in **Figures S14-18**, and summary results in **Figure 6**. Overall, liver and adipose were the most enriched and most significant tissues for lipids traits, while showing variability between exposed and unexposed individuals. LDL also showed some significance and variability for cell-types mapped to digestive tissue for the drinking exposures and current smoking. There was less significant enrichment and a less marked difference for BP traits, although we noticed a substantially larger enrichment in liver tissue among current drinkers versus non-drinkers for DBP.

**Figure 6.**
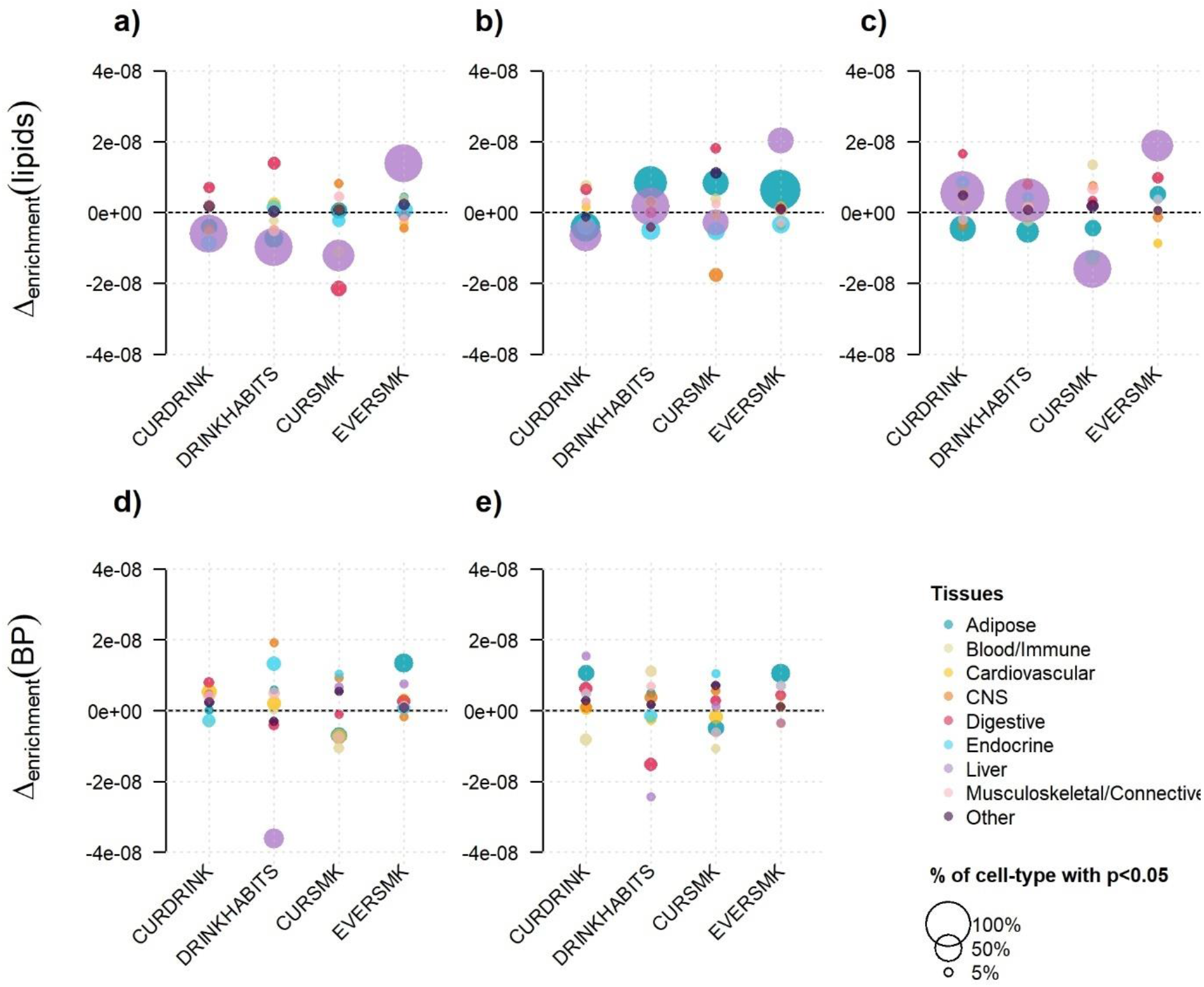
Stratification of heritability by tissue. Cell-type partitioned heritability for each exposure was performed and further merged into nine primary tissue categories. The top panels show the results for lipids: LDL (a), HDL (b), and TG (c), and the bottom panels show the results for blood pressure: DBP (d), and SBP (e). For each phenotype-exposure pair we derived the difference between the median enrichment in exposed and unexposed individuals (Δ_enrichment_) per tissue. To highlight the significance of enrichment within each cell-type, we scaled the size of each data point by the proportion of cell types that are nominally significant (i.e. P<0.05) after merging exposed and unexposed results.

### Risk of bias for the test of interaction in trans-ancestry analysis

Throughout this work, we used GxE interaction effect estimates and *p*-values derived using the standard 1df inverse-variance meta-analysis scheme as described in Willer et al^40^, and applied to the 1df interaction effect from each contributing cohort. This is similar to the approach used in the original GWIS papers. On the other hand, we used the main genetic effect estimates and *p*-values derived from the 2df framework as described in Manning et al^14^. Indeed, in addition to computing the 2df joint test *p*-value, the 2df framework provides a joint estimation of the main and interaction effect coefficients along with standard errors. These parameters can then be used to perform a *Wald* test of both the main genetic effect and the GxE interaction effect. Our choice of using only the main effect from that framework, while using the standard univariate meta-analysis for the GxE interaction, was driven by potential bias we observed in a pilot study using results from both frameworks.

Indeed, while the two frameworks (standard 1df and joint estimation) should produce asymptotically similar results, we noticed some differences at top associated variants for the interaction effect results. We therefore conducted a series of simulations exploring different scenarios to compare the performance in terms of type I error rate and power using a simple case of two cohorts. The different scenarios were designed to explore heterogeneity in allele frequencies, in proportions of exposed individuals, and in genetic and interaction effects between the two cohorts (see Material and Methods, **Table S8**). In most scenarios, *p*-values for interaction computed using the two frameworks were similar. However, we found that in the absence of an interaction effect but with heterogeneity between the two cohorts for both the main genetic effect and the proportion of exposed individuals, the test of interaction derived from the 2df framework exhibited a major increase in type I error rate (**Figure S19**). This specific pattern of both differential genetic effect and exposure frequency likely induce a confounding effect on the interaction term (i.e. higher genetic effect in a cohort with a higher level of exposure). Note that heterogeneity in the main genetic effect might be explained by a form of GxE interaction; however, it can also very likely be due to differences in linkage disequilibrium between causal and typed variants – especially when analyzing multiple ancestries – and thus inducing a false interaction signal.

### Genetic heterogeneity and power for 2df tests

To understand further the characteristics of the 2df framework, we performed an additional series of analyses using again a simple case were two cohorts with different characteristics were here pooled together. First, we performed a simulation in which we compared coefficient and chi-squared statistics for both the main genetic and the GxE interaction effect, using the 1df and 2df frameworks across P2017a large number of replicates where all parameters were drawn at random. As showed in **Figure S20**, when the exposure is binary (as in the present study), we confirm a potential bias for the interaction effect but no difference in the main genetic effect. When the exposure is continuous, we observed variability in both estimates, but no systematic bias (**Figure S21**). Second, we simulated a series of replicates and compared four models: a 1df marginal model only testing the effect of the genetic variants (Y~G), the same model but adjusting for the effect of the exposure (Y~G+E), a joint 2df model accounting for interaction between G and E, as used in our studies (Y~G+E+GxE), and an alternative joint 2df accounting for interaction between G and the cohort status (Y~G+C+GxC). As showed in **Figure S22**, under a complete null model, all tests are correctly calibrated. In the presence of homogenous genetic effect between the two cohorts, the 2df test is slightly less powerful than the marginal and adjusted model, which is expected as this test has one additional degree of freedom. However, when simulating heterogeneous genetic effetc between the two cohorts, the 2df accounting for GxE interaction shows an increase in power despite the absence of such interaction. This is likely explained by the exposure acting as a proxy for the cohorts, which itself is involved in a statistical interaction because of the heterogeneity. Indeed, the joint 2df accounting for GxCohort interaction shows the highest power in this scenario.

## Discussion

In this study, we assembled and synthesized the results from 28 gene-by-environment interaction GWIS on lipid and blood pressure phenotypes performed across four ancestries, which were recently published by the Gene-Lifestyle Interactions Working Group^5,7,8,10,11^. This analysis highlights a number of features regarding large-scale GxE analysis and trans-ancestry studies. Overall, we found the trans-ancestry 2df test to be efficient for SNP discovery, with the vast majority of associations identified in ancestry-specific analyses being confirmed in the trans-ancestry analysis, while allowing for a 10% increase in identified signals. Conversely, our data pointed toward ancestry-specific patterns for interaction effects, which might be due to differences in allelic frequencies at causal variants, but also to other unmeasured factors. For example, African-ancestry analyses displayed several interaction effects across all phenotypes-exposure pairs involving several variants almost absent in other populations. Our study also found differences when comparing results across exposures. We noted a greater increase in detection for lipid-associated variants when accounting for interaction with drinking, and a greater increase in detection for blood pressure-associated variants when accounting for interaction with smoking, thus stressing the potential importance of these phenotype-exposure pairs. Finally, our assessment of variance explained by interaction effects suggest that, even if small, accounting for interaction can help push signals above the stringent genome-wide significance threshold. Furthermore, the decomposition of heritability by functional annotations highlighted that exposures can induce divergent mechanisms of phenotype production with modification in the associated genetic pathway and cell type involved.

Comparing our results against previous GWAS of marginal genetic effect, we found strong concordance of effects for lipid analyses, with most of the previously identified loci being validated in our study, and over 190 new associations identified. Results for blood pressure were more heterogeneous, with approximately half of the known associations being validated, and as many associations being only found either in our study or in previous GWAS. These differences might potentially involve heterogeneity of genetic effect across populations, as the vast majority of the non-validated loci for DBP and SBP were found in the UK Biobank cohorts. When using the marginal genetic effect reported in these studies to select potential candidate for interaction effect, we did not observe enrichment for interaction effects among those variants, nor at other less significant variants. This is in agreement with our in-depth comparison of main genetic and interaction effects using the consortium data, which found only negligible correlation between the interaction and main effects coefficients.

Our estimation of the phenotypic variance explained by marginal genetic effect and interaction effect are in agreement with previous studies, showing that the contribution of GxE terms on top of marginal genetic effect is relatively modest. It confirms the likely limited impact of discovering GxE for prediction purposes in the general population^41^. In addition, the variability observed across population, exposures and phenotypes might be explained by various which cannot be sorted out using these data. Consequently, further work would be needed not only to understand this heterogeneity but also assess special cases, such as the prediction performances in strata defined by environmental exposure, which might lead to gain in predictive power^42^. In this regard, our exposure-specific heritability analyses suggest a potentially larger polygenic effect (and therefore a higher prediction power) among non-smokers and current drinkers for most phenotypes. Also, a modest contribution of GxE to phenotypic variance does not rule out the potentially important role of GxE in the etiology of these traits. For example, a marginal model can capture most of the variance explained by interaction effect, thus masking more complex biological mechanisms^4^. Indeed, the interaction effect only represents the deviation of genetic effect relative to the mean of the exposure. Our stratified heritability analyses provide a good example of this hypothesis, suggesting in particular a potential change in the genetic architecture of LDL conditional on smoking and BP conditional on drinking – i.e. there was much more variability in the enriched annotations between exposed and unexposed individuals for those trait-exposure pairs as compared to the other pairs.

Additionally in this study, we identified a number of methodological subtleties with the 2df framework that can make the interpretation of interaction and main effect complex, and in the worst case, can lead to false conclusions about the potential link between the two parameters and about potential interaction effect for the genetic variants under study. The detailed characterization of the method we performed provides guidelines for future studies. In particular our work highlighted the following points: i) when heterogeneity of genetic effect across the cohorts analyzed is suspected and the exposure has different distribution across those cohorts, the standard 1 degree of freedom inverse-variance meta-analysis of all cohorts should always be preferred for testing interaction effect using estimates from the joint 2df framework; ii) while the interaction effect estimate derived from the 2df framework shows reduced robustness in the presence of heterogeneity, the main effect estimate was much less sensitive to those factors in our simulation; iii) despite this potential bias in effect estimates, the joint 2df test of main genetic and interaction effect remains itself well calibrated in all scenarios we considered; and iv) we found that while genetic heterogeneity can bias the interaction test, it can at the same time boost the power of the 2df test, making it more powerful than the test of marginal genetic effect even in the absence of an interaction effect. Our simulations indicated that this gain in power is due to the exposure acting as a proxy for cohorts, which themselves display statistical interaction with the genetic variant.

The Gene-Lifestyle Interactions Working Group is a unique initiative that aims at understanding the interplay between genetics and lifestyle on human phenotypes across various ancestries. Here we present an overview of the published GxE screenings involving SNP by drinking and smoking exposures interactions on lipid and blood pressure traits using GxE summary statistics. These cross analyses identified different signals depending on the population and exposures. In addition, the summary data provided by the consortium provide opportunities for numerous additional follow-up analyses and we re-analyzed the data to get deeper insights into the biological mechanisms underlying the phenotypes conditional on the exposure. Future studies extending methodologies developed for marginal genetic effect GWAS can be used to gain further knowledge on GxE, using fine-mapping^43^, co-heritability^44^, or conditional analyses^45^ approaches.

## Acknowledgments

The various Gene-Lifestyle Interaction projects, including this summary project, are largely supported by a grant from the U.S. National Heart, Lung, and Blood Institute (NHLBI), the National Institutes of Health, R01HL118305. This work was also supported by the INCEPTION project (PIA/ANR-16-CONV-0005). This research was supported in part by the Intramural Research Program of the National Human Genome Research Institute, National Institutes of Health. Paul S. de Vries was supported by American Heart Association grant number 18CDA34110116. Yun J Sung was supported by the K25HL121091 award from NHLBI. James Gauderman was partly supported by the P01CA196569 grant from the National Institutes of Health. Full set of study-specific funding sources and acknowledgments were included in the four separate publications^7–10^

## References

1. McAllister, K. et al. Current Challenges and New Opportunities for Gene-Environment Interaction Studies of Complex Diseases. Am J Epidemiol 186, 753–761 (2017).

2. Gauderman, W.J. et al. Update on the State of the Science for Analytical Methods for Gene-Environment Interactions. Am J Epidemiol 186, 762–770 (2017).

3. Ritchie, M.D. et al. Incorporation of Biological Knowledge Into the Study of Gene-Environment Interactions. Am J Epidemiol 186, 771–777 (2017).

4. Aschard, H. A perspective on interaction effects in genetic association studies. Genet Epidemiol 40, 678–688 (2016).

5. Rao, D.C. et al. Multiancestry Study of Gene-Lifestyle Interactions for Cardiovascular Traits in 610 475 Individuals From 124 Cohorts: Design and Rationale. Circ Cardiovasc Genet 10 (2017).

6. Kraft, P., Yen, Y.C., Stram, D.O., Morrison, J. & Gauderman, W.J. Exploiting gene-environment interaction to detect genetic associations. Hum Hered 63, 111–9 (2007).

7. Feitosa, M.F. et al. Novel genetic associations for blood pressure identified via gene-alcohol interaction in up to 570K individuals across multiple ancestries. PLoS One 13, e0198166 (2018).

8. Sung, Y.J. et al. A Large-Scale Multi-ancestry Genome-wide Study Accounting for Smoking Behavior Identifies Multiple Significant Loci for Blood Pressure. Am J Hum Genet 102, 375–400 (2018).

9. Sung, Y.J. et al. A multi-ancestry genome-wide study incorporating gene-smoking interactions identifies multiple new loci for pulse pressure and mean arterial pressure. Hum Mol Genet (2019).

10. de, Vries P.S. et al. Multi-Ancestry Genome-Wide Association Study of Lipid Levels Incorporating Gene-Alcohol Interactions. Am J Epidemiol (2019).

11. Bentley, A.R. et al. Multi-ancestry genome-wide gene-smoking interaction study of 387,272 individuals identifies new loci associated with serum lipids. Nat Genet 51, 636–648 (2019).

12. Kooperberg, C. & Leblanc, M. Increasing the power of identifying gene × gene interactions in genome-wide association studies. Genet Epidemiol 32, 255–63 (2008).

13. Friedewald, W.T., Levy, R.I. & Fredrickson, D.S. Estimation of the concentration of low-density lipoprotein cholesterol in plasma, without use of the preparative ultracentrifuge. Clin Chem 18, 499–502 (1972).

14. Manning, A.K. et al. Meta-analysis of gene-environment interaction: joint estimation of SNP and SNP × environment regression coefficients. Genet Epidemiol 35, 11–8 (2011).

15. Chang, C.C. et al. Second-generation PLINK: rising to the challenge of larger and richer datasets. Gigascience 4, 7 (2015).

16. Genomes, Project C. et al. A global reference for human genetic variation. Nature 526, 68–74 (2015).

17. Kato, N. et al. Trans-ancestry genome-wide association study identifies 12 genetic loci influencing blood pressure and implicates a role for DNA methylation. Nat Genet 47, 1282–1293 (2015).

18. Liang, J. et al. Single-trait and multi-trait genome-wide association analyses identify novel loci for blood pressure in African-ancestry populations. PLoS Genet 13, e1006728 (2017).

19. Warren, H.R. et al. Genome-wide association analysis identifies novel blood pressure loci and offers biological insights into cardiovascular risk. Nat Genet 49, 403–415 (2017).

20. Below, J.E. et al. Meta-analysis of lipid-traits in Hispanics identifies novel loci, population-specific effects, and tissue-specific enrichment of eQTLs. Sci Rep 6, 19429 (2016).

21. Prins, B.P. et al. Genome-wide analysis of health-related biomarkers in the UK Household Longitudinal Study reveals novel associations. Sci Rep 7, 11008 (2017).

22. Surakka, I. et al. The impact of low-frequency and rare variants on lipid levels. Nat Genet 47, 589–97 (2015).

23. Teslovich, T.M. et al. Biological, clinical and population relevance of 95 loci for blood lipids. Nature 466, 707–13 (2010).

24. Purcell, S. et al. PLINK: a tool set for whole-genome association and population-based linkage analyses. Am J Hum Genet 81, 559–75 (2007).

25. Laville, V. et al. VarExp: estimating variance explained by genome-wide GxE summary statistics. Bioinformatics 34, 3412–3414 (2018).

26. Bulik-Sullivan, B.K. et al. LD Score regression distinguishes confounding from polygenicity in genome-wide association studies. Nat Genet 47, 291–5 (2015).

27. Laville, V. et al. Deriving stratified effects from joint models investigating Gene-Environment Interactions. bioRxiv, 693218 (2019).

28. Finucane, H.K. et al. Partitioning heritability by functional annotation using genome-wide association summary statistics. Nat Genet 47, 1228–35 (2015).

29. Lu, Q. et al. Systematic tissue-specific functional annotation of the human genome highlights immune-related DNA elements for late-onset Alzheimer’s disease. PLoS Genet 13, e1006933 (2017).

30. Finucane, H.K. et al. Heritability enrichment of specifically expressed genes identifies disease-relevant tissues and cell types. Nat Genet 50, 621–629 (2018).

31. Aschard, H., Hancock, D.B., London, S.J. & Kraft, P. Genome-wide meta-analysis of joint tests for genetic and gene-environment interaction effects. Hum Hered 70, 292–300 (2010).

32. Klarin, D. et al. Genetics of blood lipids among ~300,000 multi-ethnic participants of the Million Veteran Program. Nat Genet 50, 1514–1523 (2018).

33. Aschard, H. et al. Evidence for large-scale gene-by-smoking interaction effects on pulmonary function. Int J Epidemiol 46, 894–904 (2017).

34. Clarke, T.K. et al. Genome-wide association study of alcohol consumption and genetic overlap with other health-related traits in UK Biobank (N=112 117). Mol Psychiatry 22, 1376–1384 (2017).

35. Jorgenson, E. et al. Genetic contributors to variation in alcohol consumption vary by race/ethnicity in a large multi-ethnic genome-wide association study. Mol Psychiatry 22, 1359–1367 (2017).

36. Rasheed, H., Stamp, L.K., Dalbeth, N. & Merriman, T.R. Interaction of the GCKR and A1CF loci with alcohol consumption to influence the risk of gout. Arthritis Res Ther 19, 161 (2017).

37. Gauderman, W.J., Zhang, P., Morrison, J.L. & Lewinger, J.P. Finding novel genes by testing G × E interactions in a genome-wide association study. Genet Epidemiol 37, 603–13 (2013).

38. Zhang, P., Lewinger, J.P., Conti, D., Morrison, J.L. & Gauderman, W.J. Detecting Gene-Environment Interactions for a Quantitative Trait in a Genome-Wide Association Study. Genet Epidemiol 40, 394–403 (2016).

39. Lu, Q., Powles, R.L., Wang, Q., He, B.J. & Zhao, H. Integrative Tissue-Specific Functional Annotations in the Human Genome Provide Novel Insights on Many Complex Traits and Improve Signal Prioritization in Genome Wide Association Studies. PLoS Genet 12, e1005947 (2016).

40. Willer, C.J., Li, Y. & Abecasis, G.R. METAL: fast and efficient meta-analysis of genomewide association scans. Bioinformatics 26, 2190–1 (2010).

41. Aschard, H. et al. Inclusion of gene-gene and gene-environment interactions unlikely to dramatically improve risk prediction for complex diseases. Am J Hum Genet 90, 962–72 (2012).

42. Aschard, H., Zaitlen, N., Lindstrom, S. & Kraft, P. Variation in predictive ability of common genetic variants by established strata: the example of breast cancer and age. Epidemiology 26, 51–8 (2015).

43. Kichaev, G. et al. Integrating functional data to prioritize causal variants in statistical fine-mapping studies. PLoS Genet 10, e1004722 (2014).

44. Bulik-Sullivan, B. et al. An atlas of genetic correlations across human diseases and traits. Nat Genet 47, 1236–41 (2015).

45. Yang, J. et al. Conditional and joint multiple-SNP analysis of GWAS summary statistics identifies additional variants influencing complex traits. Nat Genet 44, 369-75, S1–3 (2012).

