## Supplementary Figures for "Large-scale multivariate multi-ancestry Interaction analyses point towards different genetic mechanisms by population and exposure"

#### Supplementary data

### Table of Contents

Supplementary Note ... 2
Joint test of interaction across multiple SNPs. ... 2
Variance explained at top SNP and heritability. ... 2
Stratified heritability ... 2
Supplementary Figures ... 4
Figure S1 QQplots ... 4
Figure S2. Proportion of exposed individuals ... 5
Figure S3. Distribution of independent signals per locus ... 6
Figure S4. Comparison of HDL results ... 7
Figure S5. Main and interaction effects among sSNPs significative with the 2df test ... 8
Figure S6. Interaction effect at previously reported loci ... 9
Figure S7. Example of power for 2 df interaction test ... 10
Figure S8. Variability in heritability estimation ... 11
Figure S9. Annotation enrichment for DBP ... 12
Figure S10. Annotation enrichment for SBP ... 13
Figure S11. Annotation enrichment for HDL ... 14
Figure S12. Annotation enrichment for LDL ... 15
Figure S13. Annotation enrichment for TG ... 16
Figure S14. Cell-type enrichment for DBP ... 17
Figure S15. Cell-type enrichment for SBP ... 18
Figure S16. Cell-type enrichment for HDL ... 19
Figure S17. Cell-type enrichment for LDL ... 20
Figure S18. Cell-type enrichment for TG ... 21
Figure S19. Example of power for 2 df interaction test ... 22
Figure S20. Robustness of the 2df framework for binary exposures ... 23
Figure S21. Robustness of the 2df framework ... 24
Figure S22. The joint test can capture genetic heterogeneity ... 25
References ... 26

### Supplementary Note

#### Joint test of interaction across multiple SNPs.

Consider a vector of $L$ single SNP interaction coefficient $\boldsymbol{S} = [\hat{\gamma}_1, \dots \hat{\gamma}_L]$ and their corresponding variance-covariance matrix $\mathbf{Q}$, which off-diagonal term equal 0, and diagonal equals to the variance of each estimate $\boldsymbol{\Gamma} = \left[\sigma^2_{\hat{\gamma}_1}, \dots \sigma^2_{\hat{\gamma}_L}\right]$. In the standard omnibus test, all interaction effects, are tested jointly, by forming the statistics $\boldsymbol{S}^{\boldsymbol{T}}\mathbf{Q}^{-1}\boldsymbol{S}$, which follows a chi-square distribution with L degree of freedom. For GRS-based interaction tests, we assume the GRS are built as the (weighted or unweighted) sum of risk alleles of the $L$ candidate SNPs. Explicitly, $uGRS = \sum_{i=1\dots L} G_i$, and $wGRS = \sum_{i=1\dots L} w_i \times G_i$, where $w_i$ is defined as marginal genetic risk estimates of the SNP. We aim at testing the significance of the $uGRS \times E$ and $wGRS \times E$. As previously demonstrated[1], the corresponding statistical tests can be respectively approximated using interaction summary statistics as $\left(\sum_{i=1\dots L}\left(\hat{\gamma}_i/\sigma^2_{\hat{\gamma}_i}\right)\right)^2/\sum_{i=1\dots L}\left(1/\sigma^2_{\hat{\gamma}_i}\right)$, and $\left(\sum_{i=1\dots L}\left(w_i\hat{\gamma}_i/\sigma^2_{\hat{\gamma}_i}\right)\right)^2/\sum_{i=1\dots L}\left(w_i^2/\sigma^2_{\hat{\gamma}_i}\right)$, and both follow a 1 degree of freedom chi-square under the null.

#### Variance explained at top SNPs.

We estimated the fraction of phenotypic variance explained by the main effects, the interaction effects and those effects jointly $f_G$, $f_I$, $f_J$ respectively using the R package *VarExp*[2]. Considering a joint regression model including interaction terms fitted in a sample of $N$ individuals, the fraction of phenotypic variance explained by the genetic main effects of a set of SNPs, their interaction effect and jointly can be estimated using only summary statistics from the joint model by $f_G = \frac{N\left({\alpha'_G}^T\Sigma^{-1}\alpha'_G\right)-q}{(N-q)var(Y)}$, $f_I = \frac{N\left({\alpha'_{INT}}^T\Sigma^{-1}\alpha'_{INT}\right)-q}{(N-q)var(Y)}$ and $f_J = \frac{N\left({\alpha'_G}^T\Sigma^{-1}\alpha'_G + {\alpha'_{INT}}^T\Sigma^{-1}\alpha'_{INT}\right)-q}{(N-q)var(Y)}$ respectively; where $\alpha'_G$ and $\alpha'_{INT}$ denote the standardized main genetic and interaction effects and $q$ is the rank of the SNP correlation matrix $\Sigma$.

#### Heritability

We computed the genetic heritability from the summary statistics of the different screenings using the LD Score. Briefly, regressing the Wald test statistics ($\chi^2$) against the LDscores ($l$) which quantify the amount of linkage disequilibrium of a SNP with other variants provides an estimate of the genetic heritability $h^2$ as $\mathbb{E}\left[\chi^2_j | l_j\right] = Nh^2 l_j/M + Na + 1$, where $M$ is the number of SNPs, $N$ is the sample size and $a$ quantifies the contribution of confounding factors. A metric indicating the validity of the computation is the ration between the intercept of the regression and the mean Chi-square. This ratio measures the proportion of the inflation in the mean $\chi^2$ that the intercept from the regression model ascribes to causes other than polygenic heritability. This ratio is expected to be close to 0 while values substantially deviating from 0 indicate likely poor estimations of the genetic heritability due to a potential mismatch between the sample and the reference population or model specifications. In our studies, we observed some ratios largely deviating from 0 so we conducted a sensitivity analysis by estimating the genetic heritability after filtering SNPs in our study based on their p-values for the test of heterogeneity across cohorts or population. In practice, we kept SNPs with a p-value higher than a given threshold (P>0, >0.3, >0.5, >0.8, >0.9) and computed the genetic using this set of SNPs. Finally, we retained the estimation from the set of SNPs leading to the smallest ratio and a number of included SNPs greater than 200,000.

#### Stratified heritability

We used two distinct sets of annotations: *baseline* and *Genoskyline+.* The baseline annotations encompass 53 tissue-agnostic, general functional annotations. Annotations in this set are binary and are not mutually exclusive; each variant is labeled with an indicator variable for each annotation and may belong to multiple annotation classes. GenoSkyline+ is a recently proposed annotation set integrating a rich collection of epigenomic data from the Roadmap Epigenomics Project[3]. The suite of annotations are tissue-specific, genome-wide predictions of functionality, generated through integrative analysis of epigenetic and gene expression. Unsupervised learning methods were used to generate posterior probabilities of functionality at every genomic coordinate within the human genome for 127 tissue-specific annotations. The authors provide pregenerated data for use with the LDscore software; for each Hapmap3 variant, each annotation is dichotomized by assigning a positive indicator if the posterior probability is greater than or equal to 0.5, and a negative indicator if the posterior is less than 0.5.

#### Supplementary Figures

##### Figure S1 QQplots

QQplots for 1df tests of interaction for the four exposures: current drinking (a), drinking habits (b), current smoking (c), and ever smoking (d), and the 2df tests of main and interaction effect: current drinking (e), drinking habits (f), current smoking (g), and ever smoking (h).

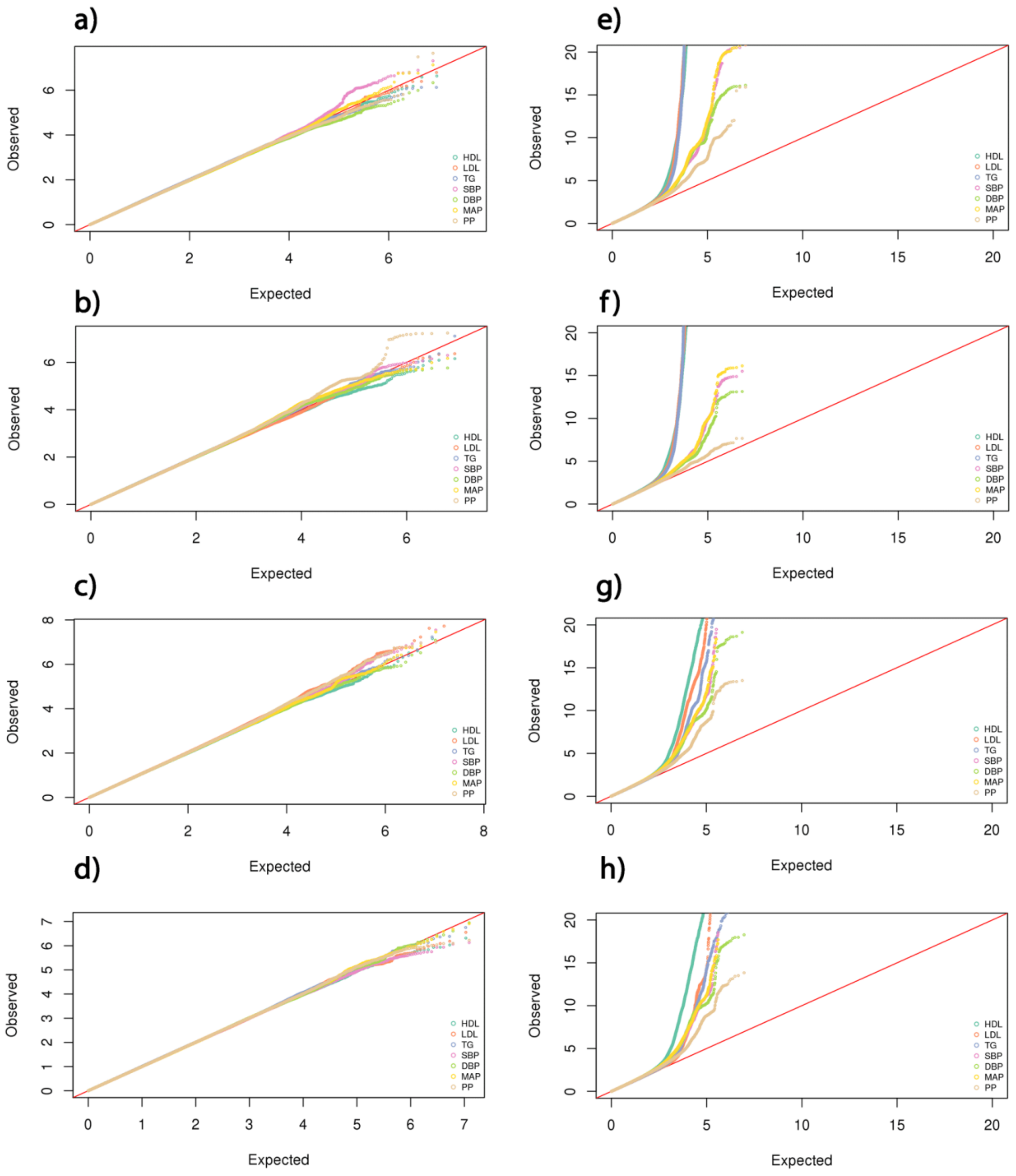

#### Figure S2. Proportion of exposed individuals

The plots show the proportion of exposed individuals (in %) per ancestry and phenotype, for each exposure.

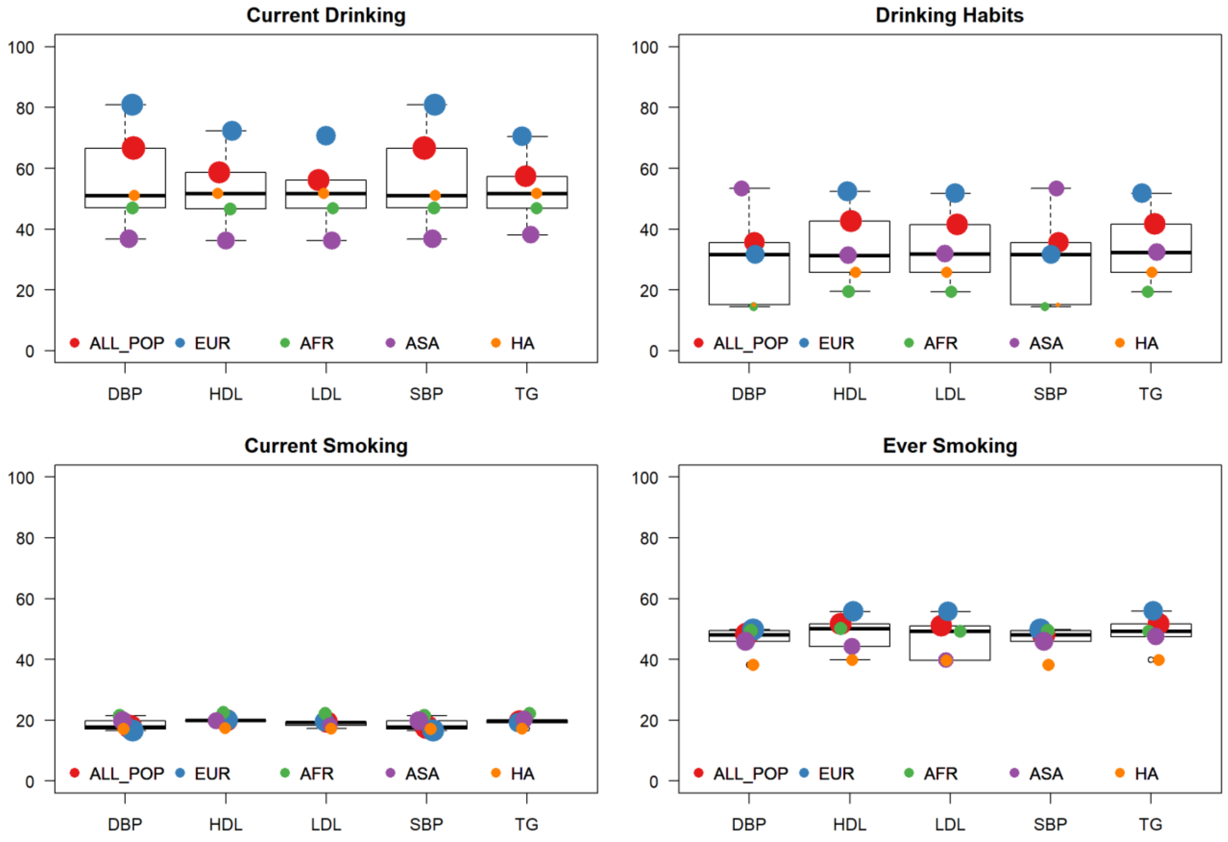

#### Figure S3. Distribution of independent signals per locus

We derived the distribution of the number of independent association signal per locus for the 2df joint test for the trans-ancestry analysis (ALL_POP), and each ancestry: European (EA), Asian (ASA), African (AA), and Hispanic (HA). Distribution were derived after merging results from all exposure-phenotypes pairs. Blue bars indicate the count of loci for each category. Red lines show the cumulative number of loci.

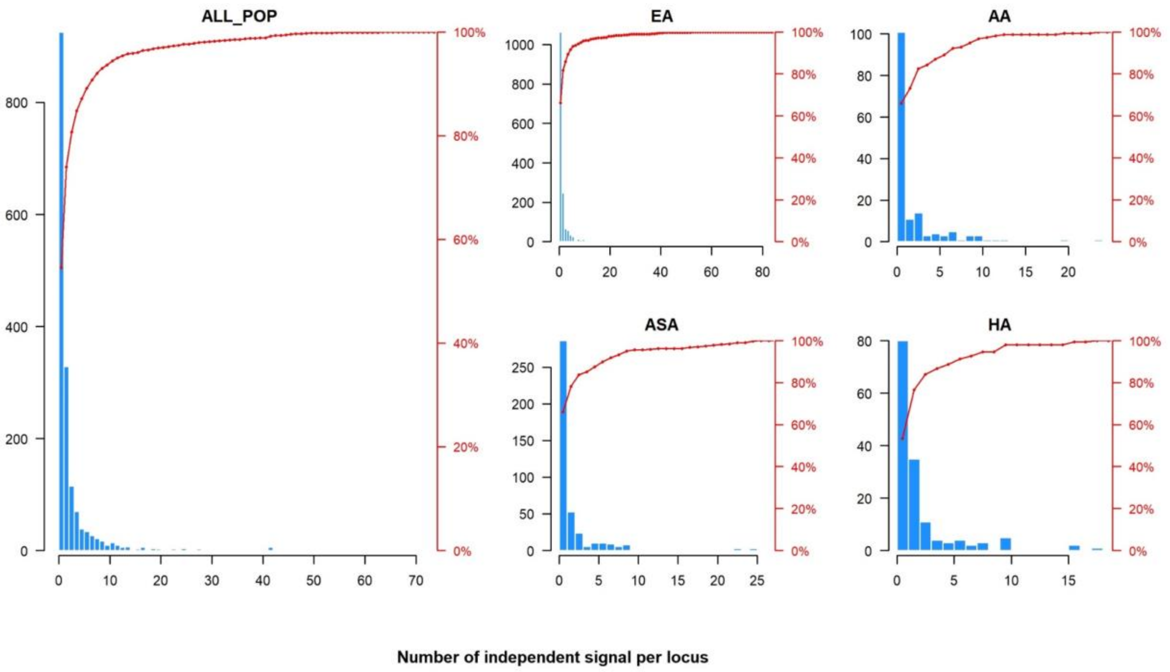

#### Figure S4. Comparison of HDL results

We compared the identified loci for HDL across the four exposures. Panel (a) shows the 2DF chi-squared at loci identified by the SNP by current drinking interaction GWAS (x axis) that were also identified by the other exposures GWAS (y axis). For those loci, the signal was highly correlated as expected. Panel (b) shows the mean chi-squared for interaction effect for the 108, 103, 67, and 69 loci found by the current drinking, drinking habits, current smoking and ever smoking exposures, respectively. Grey bars correspond to the chi-squared derived using the standard 1df METAL, and pink bars correspond to the chi-squared derived using the estimates from the 2df METAL.

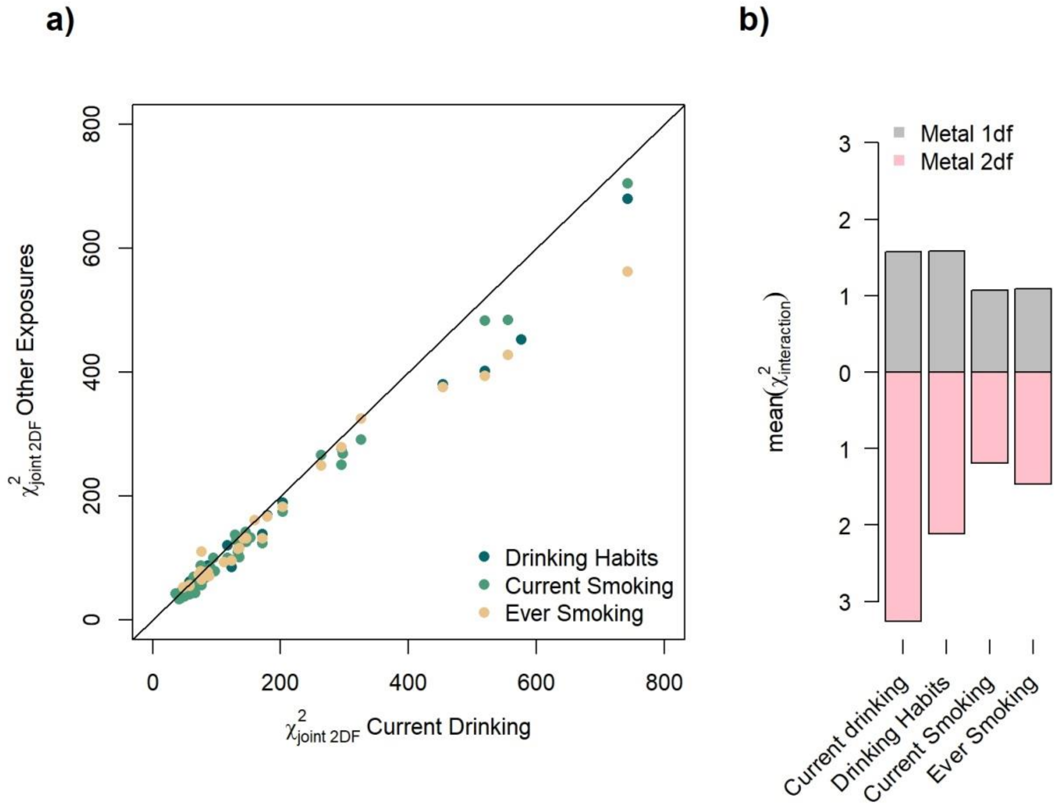

**Figure S5. Main and interaction effects among sSNPs significative with the 2df test**

For each phenotype-exposure-ancestry trio we extracted the independent SNPs found associated with the 2df test. We then derived the number of SNPs for which main and interaction coefficient were concordant (both positive or both negative, grey bar) and discordant (orange + red bar). The former pattern indicates enhanced genetic effect among exposed individuals, while the latter indicates decreased genetic effect among exposed individuals. Across SNPs with discordant main and interaction effect, the red bars show the few cases were the discordant interaction effect is large enough to induce an opposite genetic effect between exposed and unexposed.

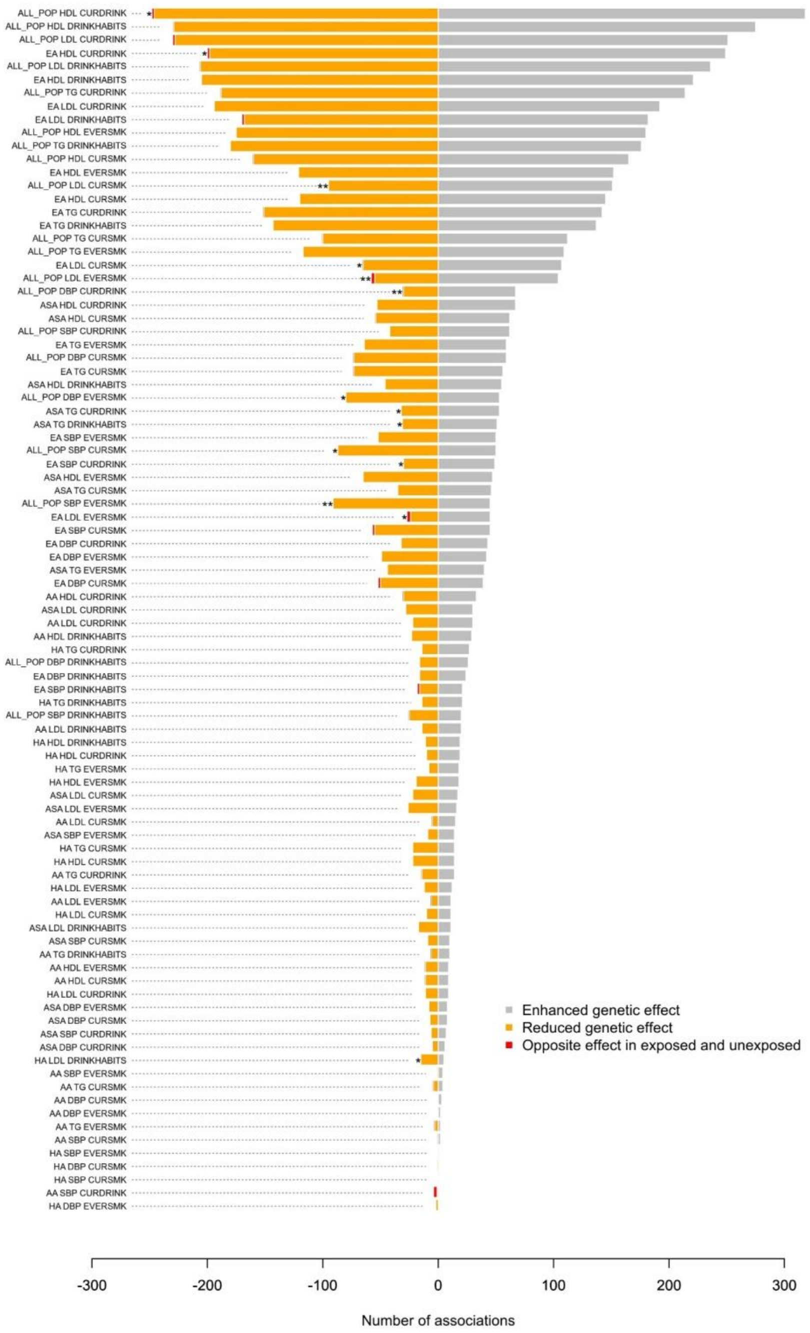

#### Figure S6. Interaction effect at previously reported loci

Forest plot of single SNP interaction Z-scores for independent genome-wide significant SNPs extracted from previous large-scale GWAS. For each SNP, we re-derived Z-score so that the coded allele is the allele associated with increasing phenotypic values in the previous marginal GWAS. Z-scores for interaction were plotted separately for current drinking (A), drinking habits (B), current smoking (C), and ever smoking (D), along the Z-scores for the marginal model (E).

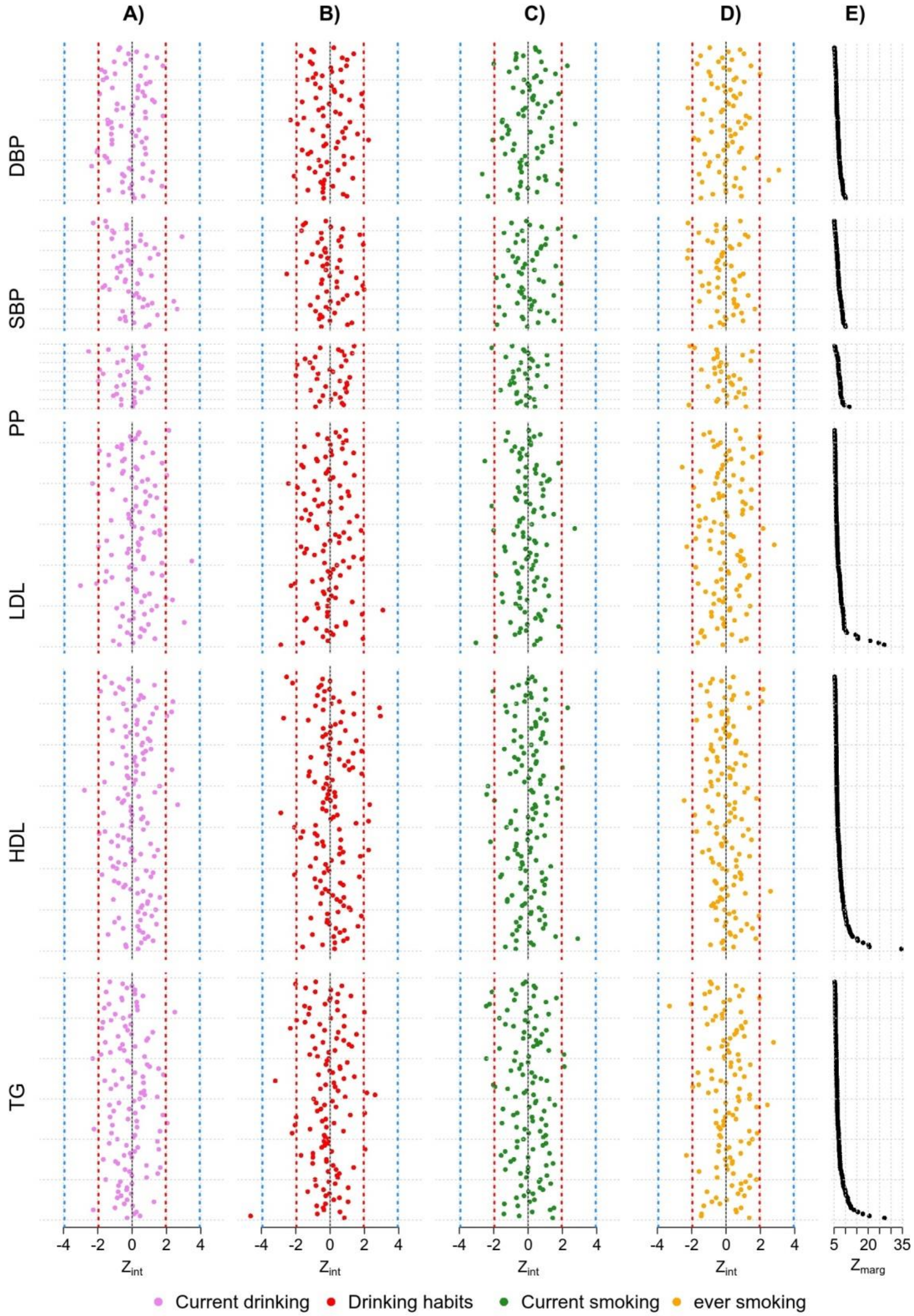

#### Figure S7. Example of power for 2 df interaction test

The following assumes a GWAS of 1 million SNPs (per SNP type I error 5e-8), N=10,000 subjects, allele freq qA=0.25 with additive genetic model, binary exposure with prevalence pE=0.4, Bg=Be=0 (no main effects), population standard deviation of Y = 1.0. The following shows power to detect GxE interaction for a range of interaction effect (*GxE Effect Size*), for the standard 1-df GxE test, 2-df joint G,GxE test, and the 2-step procedure screening on G in Step 1 and testing GxE in Step 2. Calculations were performed using the software package Quanto+[4].

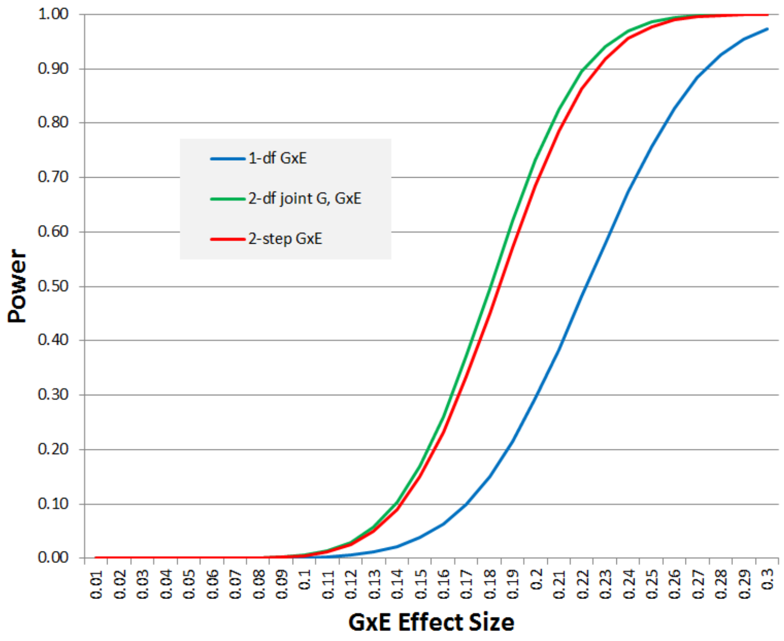

#### Figure S8. Variability in heritability estimation

We estimated heritability for each phenotype in exposed and unexposed individuals separately using LDSC, while filtering out an increasing number of variants based on their p-value for heterogeneity in the meta-analysis (P>0, >0.3, >0.5, >0.8, >0.9). Panel (a) shows the estimated heritabilities as a function of the ratio of the LDSC intercept-1 over the mean chi-squared-1. Panel (b) shows the estimated heritabilities as a function of the number of SNPs used for the estimation. Panel (c) shows the heritability per P-value threshold for each phenotype-exposure status pair. Among those, the baseline heritability (estimated without filtering, i.e. for P>0) is indicated by a green circle, and the selected heritability (with the lowest ratio and a number of SNPs larger than 200K) is indicated by a red dot.

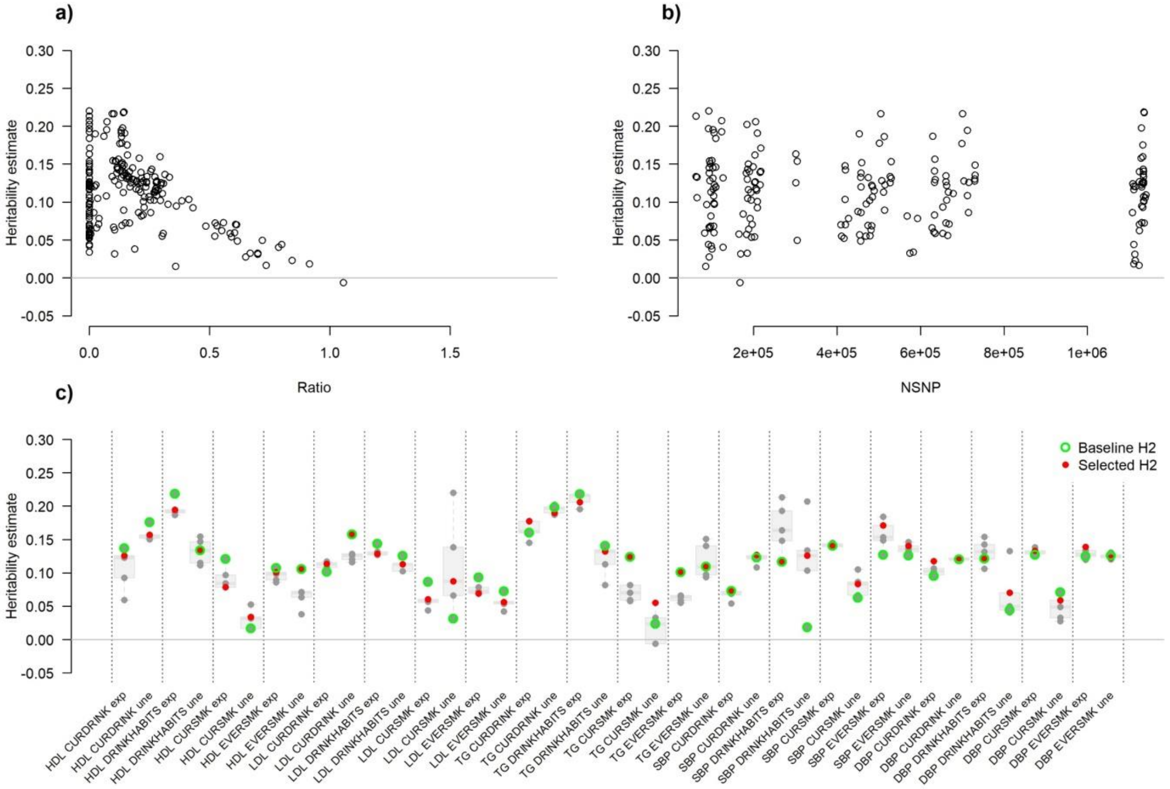

#### **Figure S9. Annotation enrichment for DBP**

Partitioned heritability for DBP in European ancestry individuals. Summary statistics from combined and stratified analyses ("Marginal": combined exposure groups for current smoking (largest sample on average), "+": exposed, "-": unexposed) were used. Color indicates enrichment of heritability. All annotations significant after correction for the 185 tests performed (P < 0.00027) are indicated by the "*" sign. For clarity, we only showed annotations significant in at least one exposure stratum.

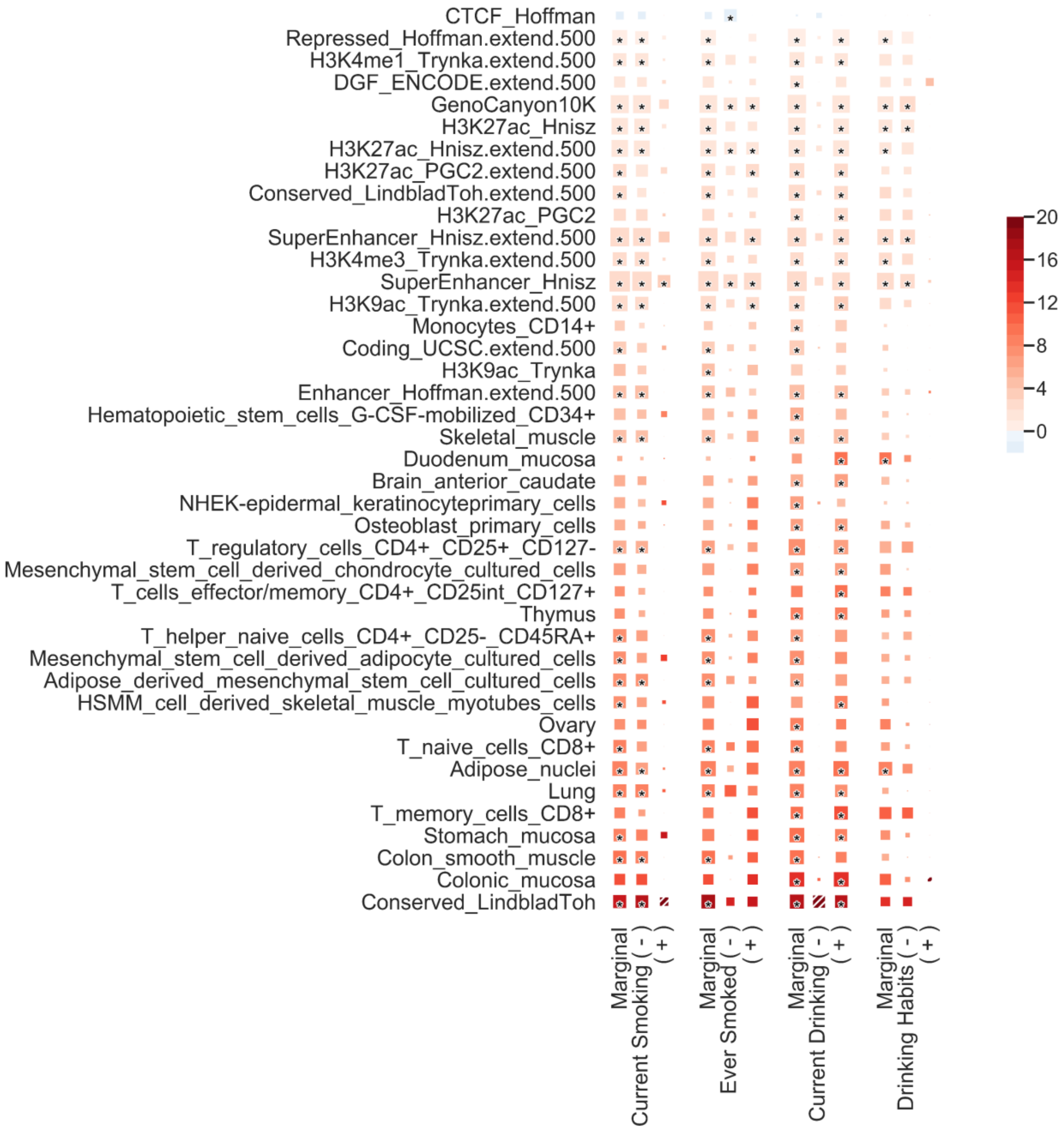

**Figure S10. Annotation enrichment for SBP**

Partitioned heritability for SBP in European ancestry individuals. Summary statistics from combined and stratified analyses (“Marginal”: combined exposure groups for current smoking (largest sample on average), "+": exposed, "-": unexposed) were used. Color indicates enrichment of heritability. All annotations significant after correction for the 185 tests performed (P < 0.00027) are indicated by the "*" sign. For clarity, we only showed annotations significant in at least one exposure stratum.

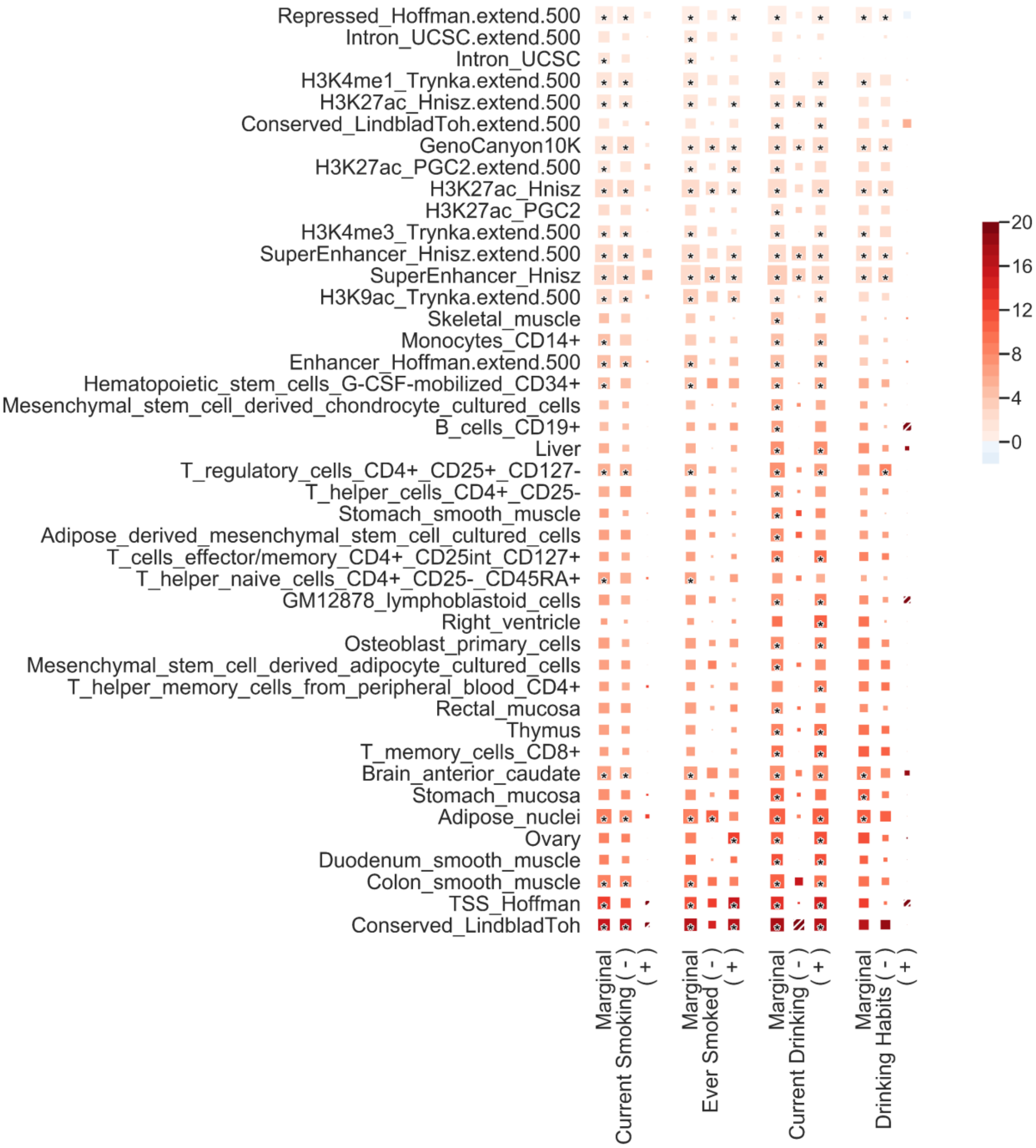

#### Figure S11. Annotation enrichment for HDL

Partitioned heritability for HDL in European ancestry individuals. Summary statistics from combined and stratified analyses (“Marginal”: combined exposure groups for current smoking (largest sample on average), "+": exposed, "-": unexposed) were used. Color indicates enrichment of heritability. All annotations significant after correction for the 185 tests performed (P < 0.00027) are indicated by the "*" sign. For clarity, we only showed annotations significant in at least one exposure stratum.

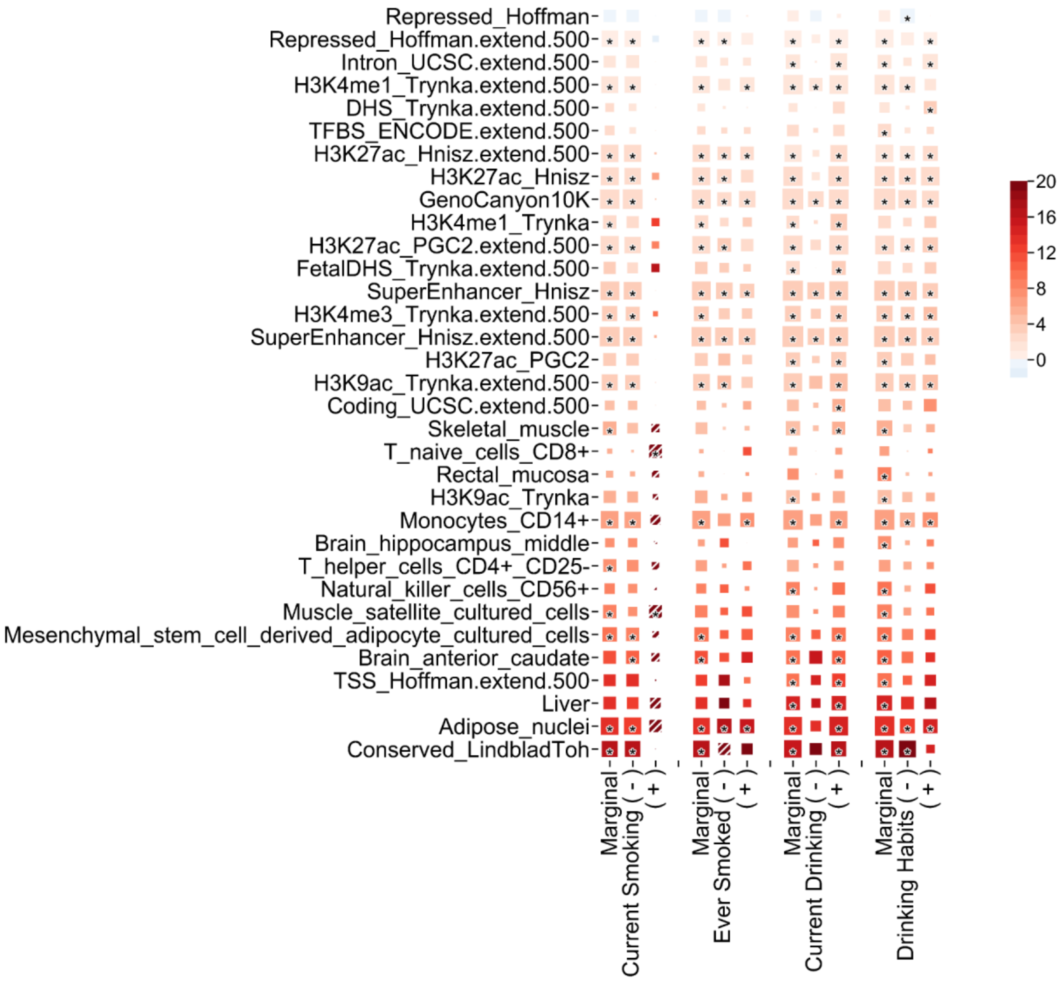

**Figure S12. Annotation enrichment for LDL**

Partitioned heritability for LDL in European ancestry individuals. Summary statistics from combined and stratified analyses (“Marginal”: combined exposure groups for current smoking (largest sample on average), "+": exposed, "-": unexposed) were used. Color indicates enrichment of heritability. All annotations significant after correction for the 185 tests performed (P < 0.00027) are indicated by the "*" sign. For clarity, we only showed annotations significant in at least one exposure stratum.

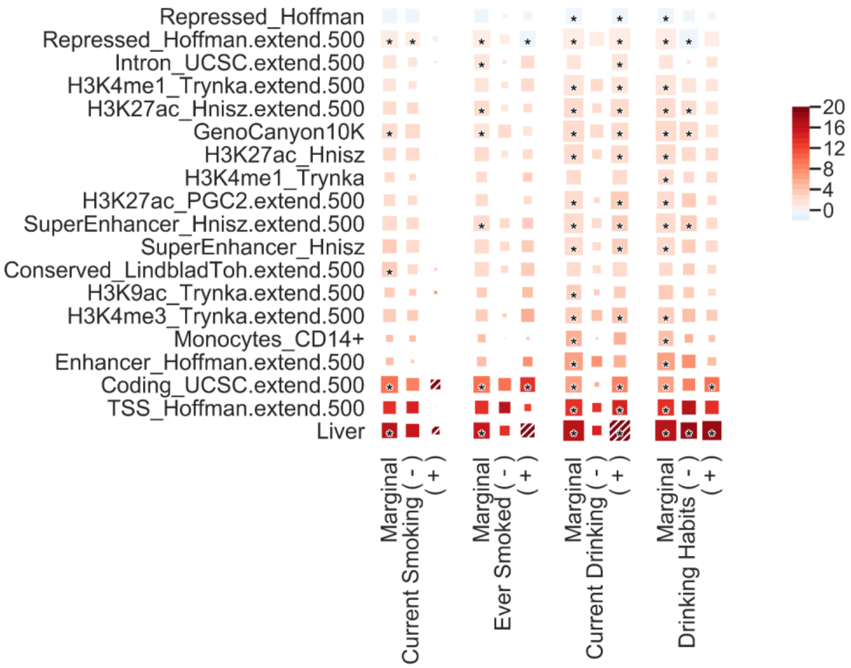

**Figure S13. Annotation enrichment for TG**

Partitioned heritability for TG in European ancestry individuals. Summary statistics from combined and stratified analyses ("Marginal": combined exposure groups for current smoking (largest sample on average), "+": exposed, "-": unexposed) were used. Color indicates enrichment of heritability. All annotations significant after correction for the 185 tests performed (P < 0.00027) are indicated by the "*" sign. For clarity, we only showed annotations significant in at least one exposure stratum.

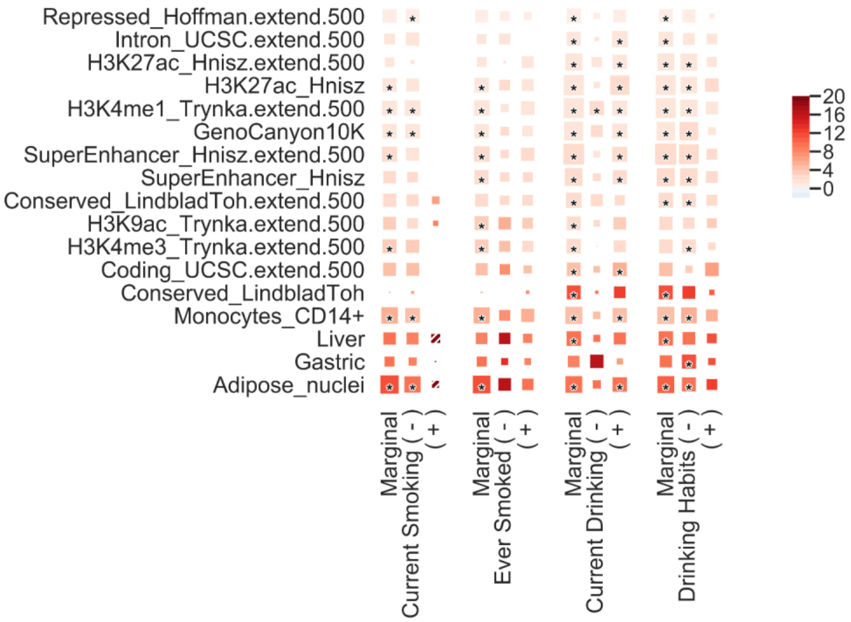

**Figure S14. Cell-type enrichment for DBP**

Cell-type partitioned of DBP heritability for each exposure: current drinking (a), current smoking (b), drinking habits (c), and ever smoking (d). Each panel shows the enrichment for the 205 cell-types on the Y axis, aggregated into tissues. For each tissue we further derived the overall median enrichment (bold line). Tissues are ordered by the median derived in unexposed individuals.

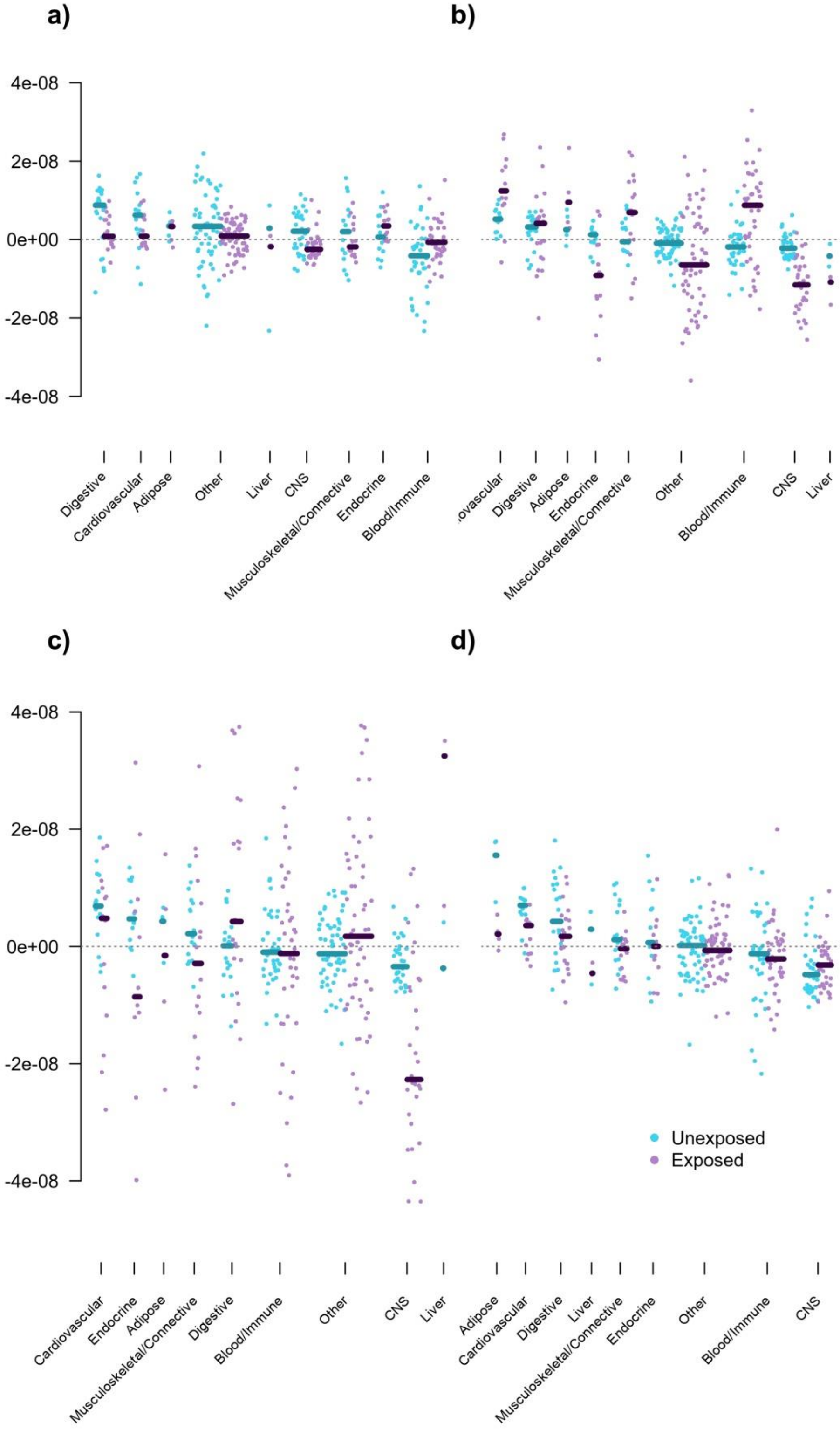

**Figure S15. Cell-type enrichment for SBP**

Cell-type partitioned of SBP heritability for each exposure: current drinking (a), current smoking (b), drinking habits (c), and ever smoking (d). Each panel shows the enrichment for the 205 cell-types on the Y axis, aggregated into tissues. For each tissue we further derived the overall median enrichment (bold line). Tissues are ordered by the median derived in unexposed individuals.

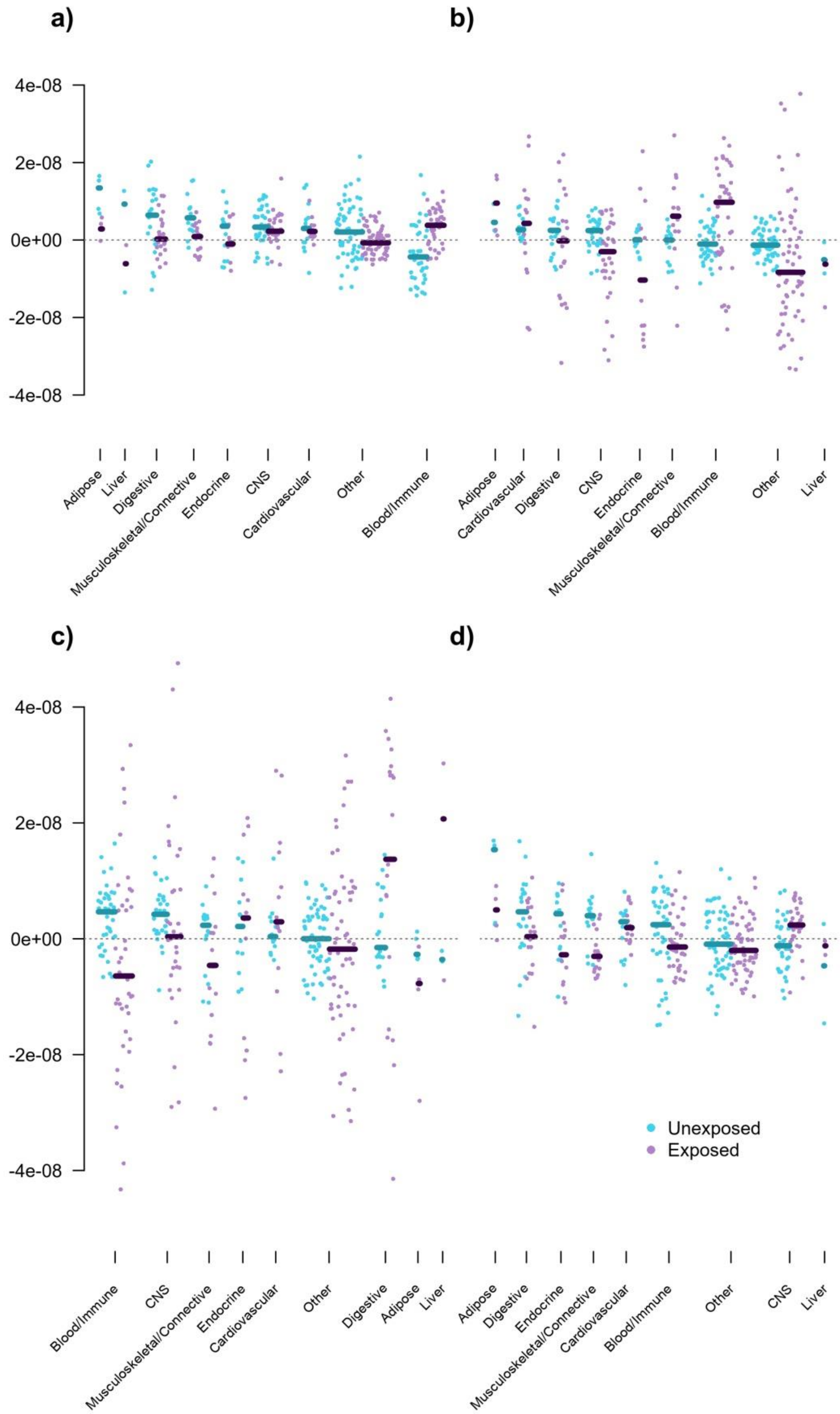

#### Figure S16. Cell-type enrichment for HDL

Cell-type partitioned of HBP heritability for each exposure: current drinking (a), current smoking (b), drinking habits (c), and ever smoking (d). Each panel shows the enrichment for the 205 cell-types on the Y axis, aggregated into tissues. For each tissue we further derived the overall median enrichment (bold line). Tissues are ordered by the median derived in unexposed individuals.

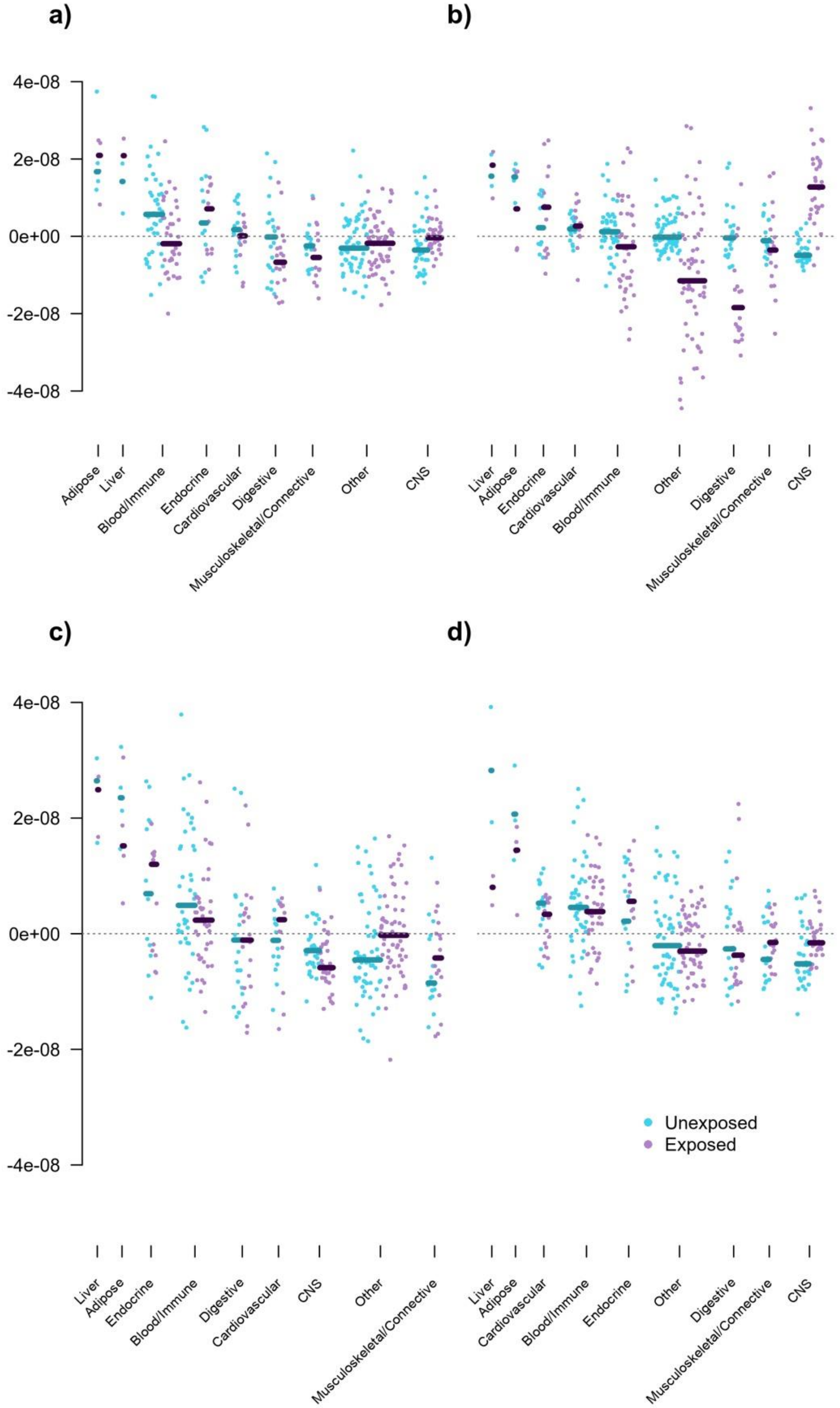

**Figure S17. Cell-type enrichment for LDL**

Cell-type partitioned of LDL heritability for each exposure: current drinking (a), current smoking (b), drinking habits (c), and ever smoking (d). Each panel shows the enrichment for the 205 cell-types on the Y axis, aggregated into tissues. For each tissue we further derived the overall median enrichment (bold line). Tissues are ordered by the median derived in unexposed individuals.

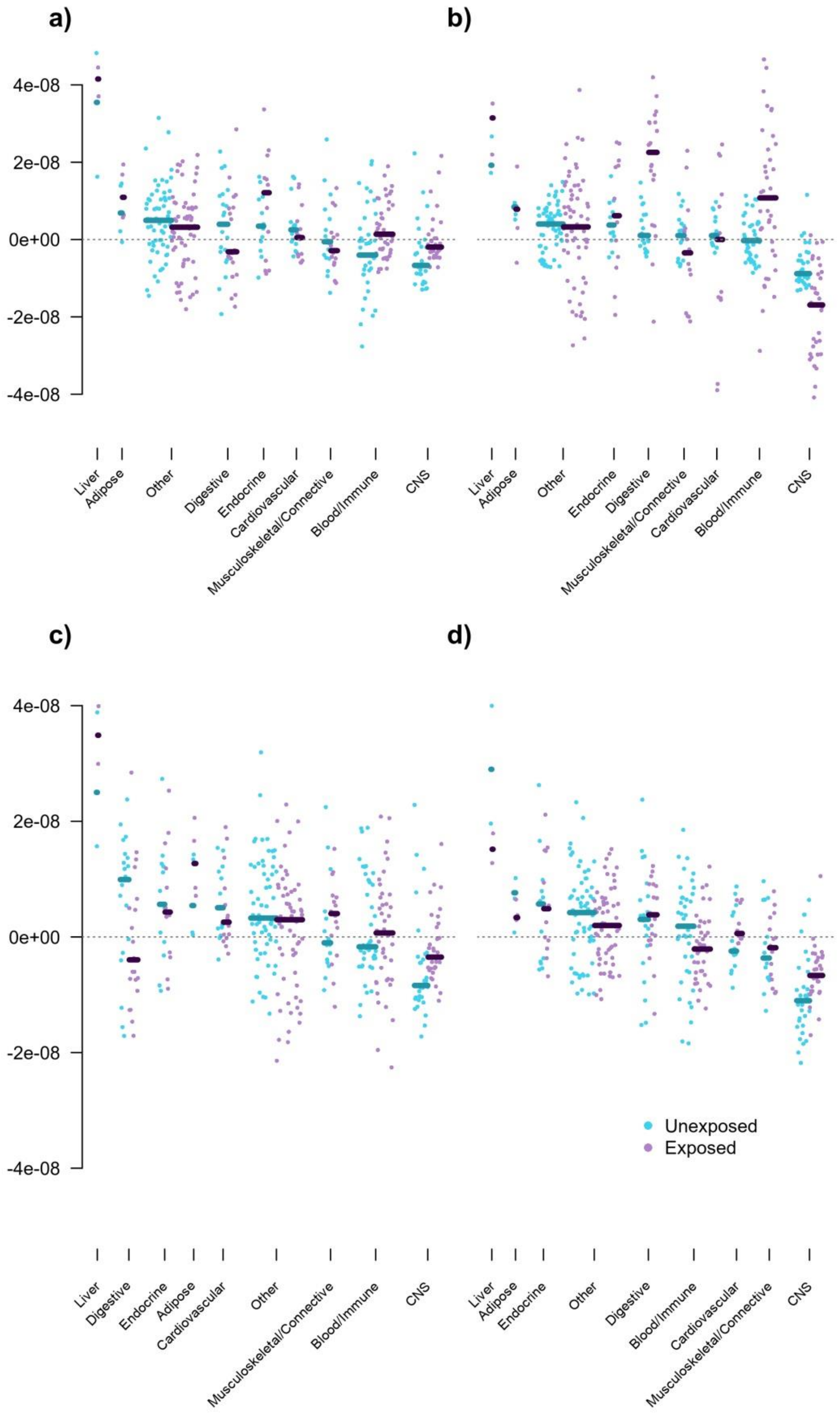

#### Figure S18. Cell-type enrichment for TG

Cell-type partitioned of TG heritability for each exposure: current drinking (a), current smoking (b), drinking habits (c), and ever smoking (d). Each panel shows the enrichment for the 205 cell-types on the Y axis, aggregated into tissues. For each tissue we further derived the overall median enrichment (bold line). Tissues are ordered by the median derived in unexposed individuals.

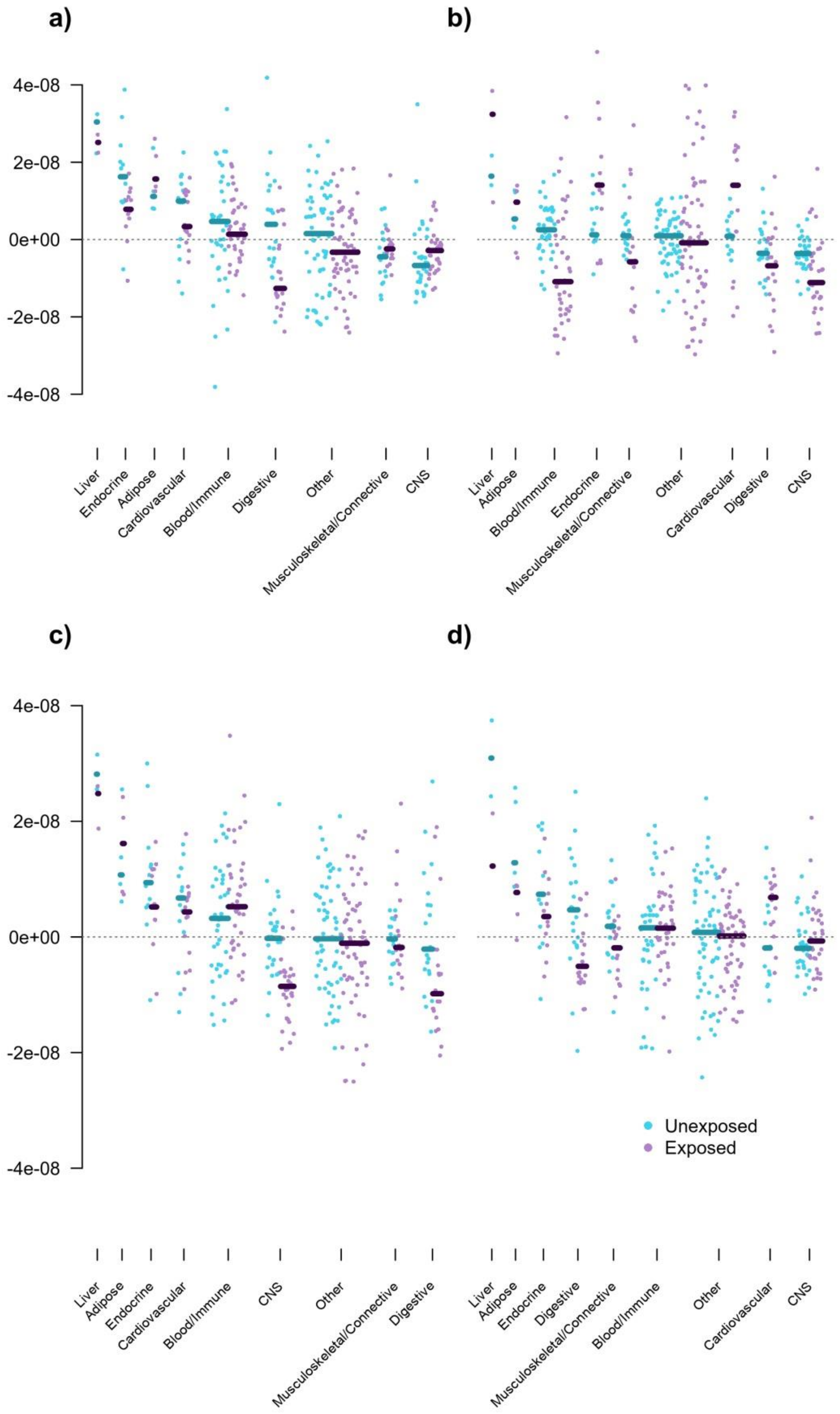

**Figure S19. Example of power for 2 df interaction test**

We simulated genotype data in two cohorts of 20,000 individuals under different scenarios to explore the impact of heterogeneity across cohorts (**Table S7**).We assessed the significance of the interaction effect in each cohort and by performing a meta-analysis using either the inverse-variance weighting scheme or the 2 degrees of freedom joint framework. We plotted the p-values distribution for the two cohorts and the two methods for each scenario.

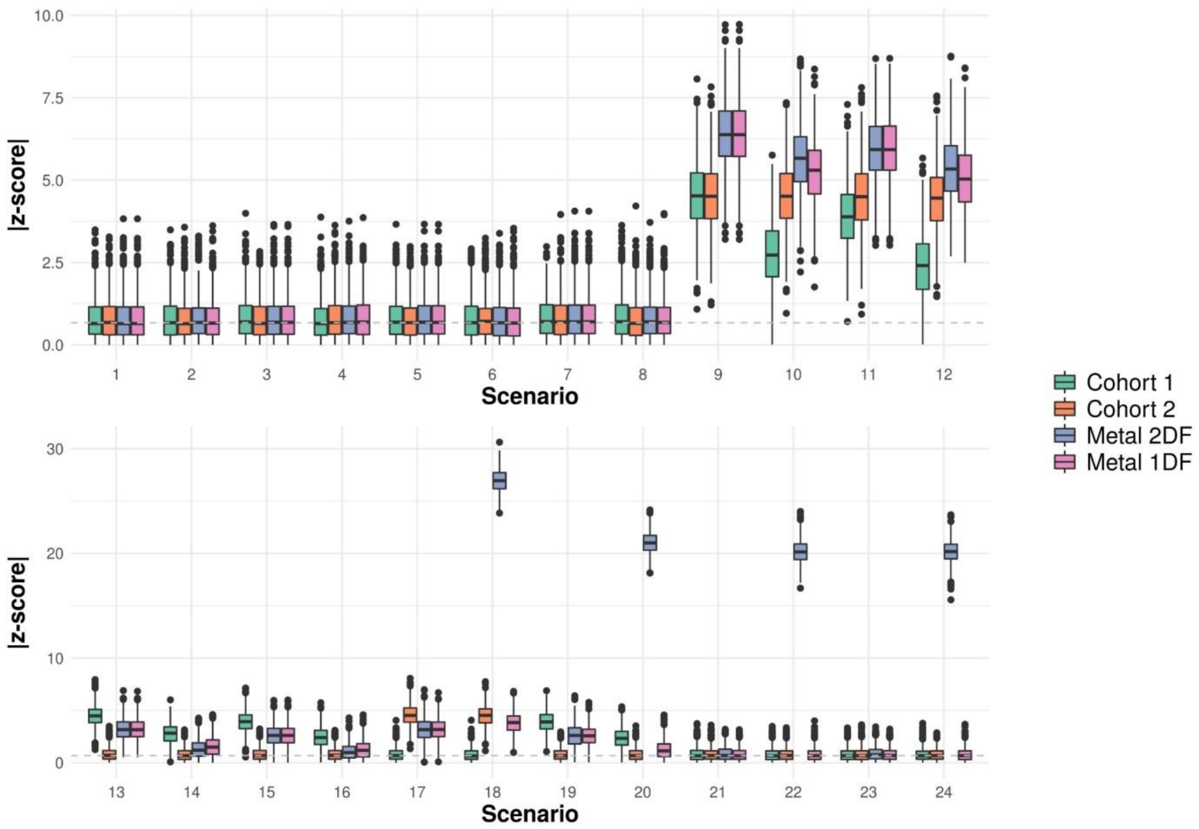

#### Figure S20. Robustness of the 2df framework for binary exposures

We simulated series of 1,000 replicates, each of them including data on a single genotype (G), a binary exposure (E) and a continuous outcome (Y) for two independent cohorts. For each replicate we performed a linear regression for each cohort using the model Y=E+G+GxE. We then derived the effect estimates and the associated chi-squared of both the main genetic effect and the interaction effect using a standard inverse-variance meta-analysis of the two cohort (1df), and using the 2df framework. We randomly varied all simualtion parameters across replicates and independently in each cohort, including the sample size per cohort (from 5,000 to 15,000), the genotype frequency (from 0.01 to 0.99), the frequency of the exposure, and including or not main and interaction effects in either cohort. Left panel shows the estimated main (*beta_main*) and interaction (*beta_int*) effects for the 1df (X axis) and the 2df (Y axis) approaches, respectively. Right panels show the chi-squared statistics for the corresponding analysis. Data points in red correspond to null model where the generative model does not include main (top panels) or interaction (bottom panels) effects.

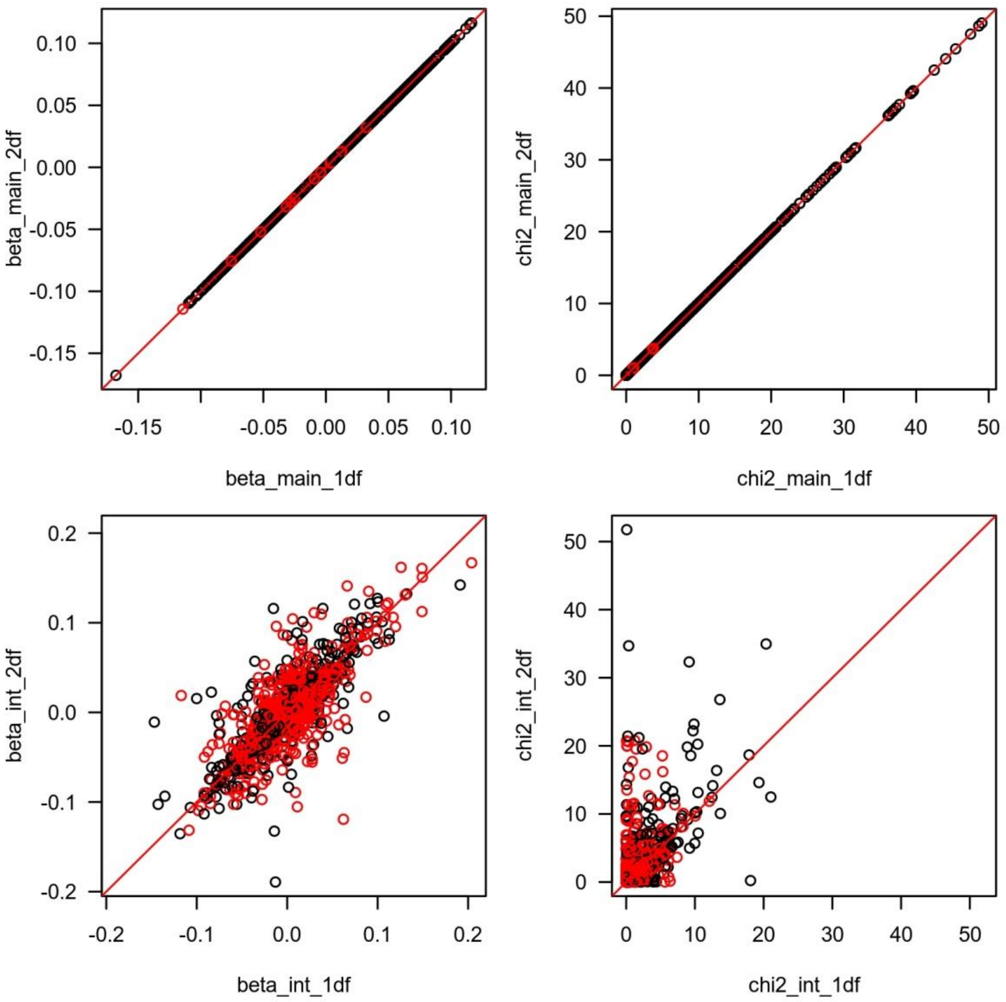

#### Figure S21. Robustness of the 2df framework

We simulated series of 1,000 replicates, each of them including data on a single genotype (G), a continuous exposure (E) and a continuous outcome (Y) for two independent cohorts. For each replicate we performed a linear regression for each cohort using the model Y=E+G+GxE. We then derived the effect estimates and the associated chi-squared of both the main genetic effect and the interaction effect using a standard inverse-variance meta-analysis of the two cohort (1df), and using the 2df framework. We randomly varied all simualtion parameters across replicates and independently in each cohort, including the sample size per cohort (from 5,000 to 15,000), the genotype frequency (from 0.01 to 0.99), the mean and variance of the exposure, and including or not main and interaction effects in either cohort. Left panel shows the estimated main (*beta_main*) and interaction (*beta_int*) effects for the 1df (X axis) and the 2df (Y axis) approaches, respectively. Right panels show the chi-squared statistics for the corresponding analysis. Data points in red correspond to null model where the generative model does not include main (top panels) or interaction (bottom panels) effects.

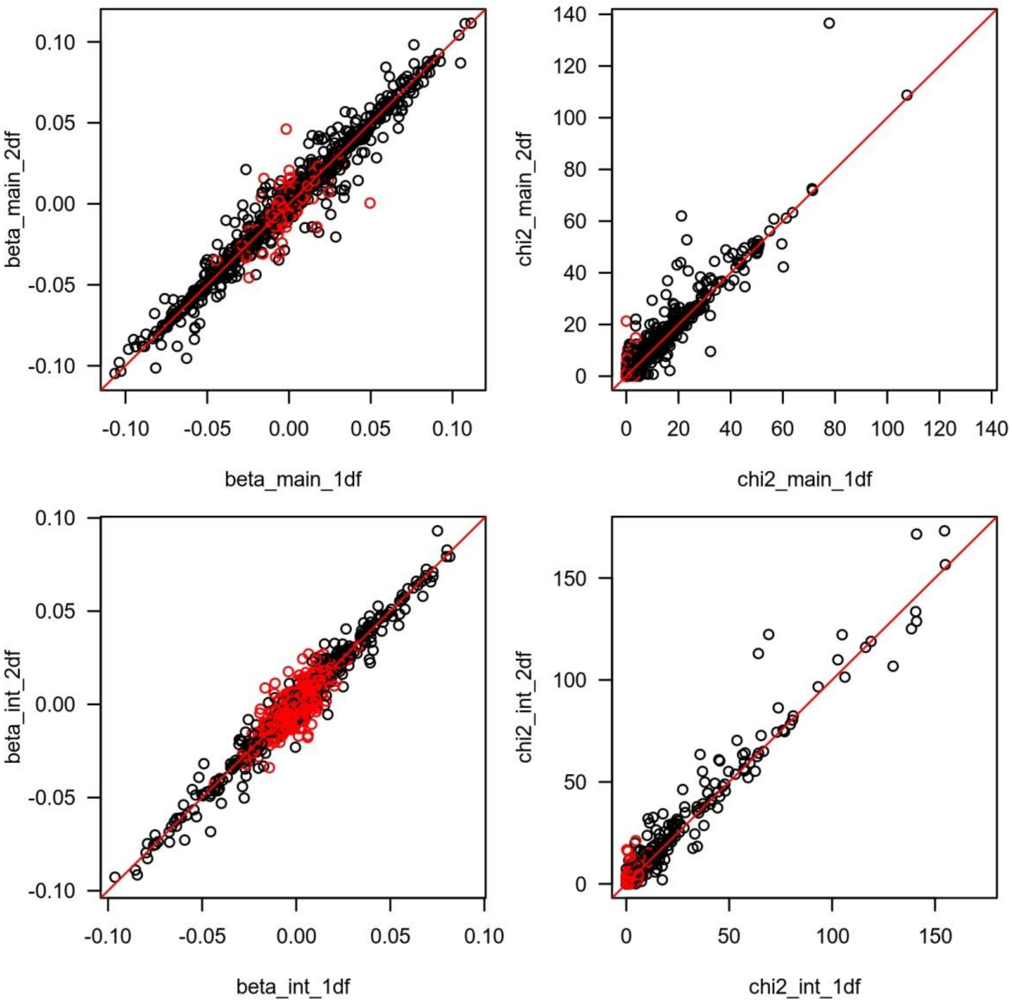

**Figure S22. The joint test can capture genetic heterogeneity**

We simulated series of 1,000 replicates, each of them including data on a single genotype (G), a binary exposure (E) and a continuous outcome (Y) for two independent cohorts. For each replicate we pooled the two cohorts and performed a linear regression for each cohort using four models: a 1df marginal model only testing the effect of the genetic variants (Y~G), the same model but adjusting for the effect of the exposure (Y~G+E), a joint 2 df model accounting for interaction between G and E, as used in our studies (Y~G+E+GxE), and an alternative joint 2 df accounting for interaction between G and the cohort status (Y~G+C+GxC). The three panels show the QQplot for the four tests under three generative scenarios. Panel (a) is a null model where there is no genetic effect in neither cohort. Panel (b) is a parsimonuious model where there is homogeneous genetic effect in both cohort. Finally, in panel (c), data were generated assuming heterogeneity between the two cohorts, with some genetic effect only in one cohort.

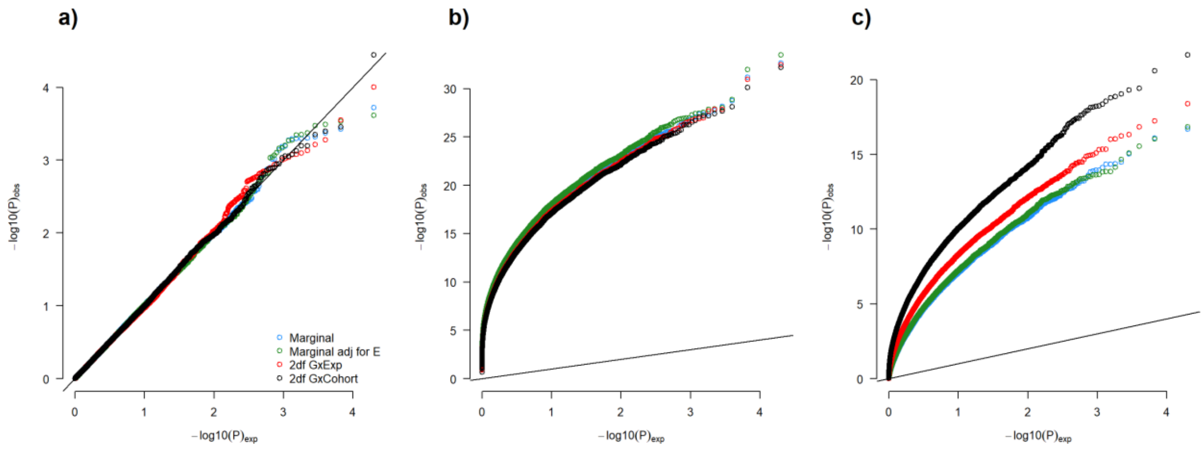

### References


1. Aschard, H. A perspective on interaction effects in genetic association studies. *Genet Epidemiol* **40**, 678-688 (2016).
2. Laville, V. *et al.* VarExp: estimating variance explained by genome-wide GxE summary statistics. *Bioinformatics* **34**, 3412-3414 (2018).
3. Lu, Q. *et al.* Systematic tissue-specific functional annotation of the human genome highlights immune-related DNA elements for late-onset Alzheimer's disease. *PLoS Genet* **13**, e1006933 (2017).
4. Murcray, C.E., Lewinger, J.P., Conti, D.V., Thomas, D.C. & Gauderman, W.J. Sample size requirements to detect gene-environment interactions in genome-wide association studies. *Genet Epidemiol* **35**, 201-10 (2011).
